# Unique spatiotemporal fMRI dynamics in the awake mouse brain

**DOI:** 10.1101/2021.08.24.457302

**Authors:** Neha Atulkumar Singh, Daniel Gutierrez-Barragan, Filomena Alvino, Ludovico Coletta, Federico Rocchi, Elizabeth De Guzman, Alberto Galbusera, Stefano Panzeri, Mauro Uboldi, Alessandro Gozzi

## Abstract

Human imaging studies have shown that spontaneous brain activity exhibits stereotypic spatiotemporal reorganization in awake, conscious conditions with respect to minimally conscious states. However, whether and how this phenomenon can be generalized to lower mammalian species, remains unclear. Leveraging a robust protocol for resting-state fMRI (rsfMRI) mapping in non-anesthetized, head-fixed mice, we investigated functional network topography and dynamic structure of spontaneous brain activity in wakeful animals. We found that rsfMRI networks in the awake state, while anatomically comparable to those observed under anesthesia, are topologically configured to maximize interregional communication, departing from the underlying community structure of the mouse axonal connectome. We further report that rsfMRI activity in wakeful animals exhibits unique spatiotemporal dynamics characterized by a state-dependent, dominant occurrence of coactivation patterns encompassing a prominent participation of arousal-related forebrain nuclei, and functional anti-coordination between visual-auditory and polymodal cortical areas. We finally show that rsfMRI dynamics in awake mice exhibits a stereotypical temporal structure, in which state-dominant coactivation patterns are configured as network attractors. These findings suggest that spontaneous brain activity in awake mice is critically shaped by state-specific involvement of basal forebrain arousal systems, and document that its dynamic structure recapitulates distinctive, evolutionarily-relevant principles that are predictive of conscious states in higher mammalian species.

## Introduction

Recent years have seen an increased interest in the application of resting state fMRI (rsfMRI) in physiologically accessible species (Gozzi and Schwarz, 2016; Milham et al., 2020). The use of these methods in the mouse has highlighted encouraging cross-species correspondences in the organization of functional networks (Gozzi and Schwarz, 2016; Whitesell et al., 2021), offering novel opportunities to mechanistically probe the neural basis of brain-wide fMRI coupling and its breakdown in brain disorders (Bertero et al., 2018; Canella et al., 2020; Pagani et al., 2020).

To ensure immobilization of animals during image acquisition and prevent motion-related artefacts, the vast majority of mouse rsfMRI studies to date have been carried out using light anesthesia (Sforazzini et al., 2014; Grandjean et al., 2020). While the employed protocols have been shown to preserve the functional (Sforazzini et al., 2014; Coletta et al., 2020) and dynamic architecture (Gutierrez-Barragan et al., 2019) of rsfMRI networks in this species, anesthetic agents can alter hemodynamic and neurovascular coupling (Rungta et al., 2017; Slupe and Kirsch, 2018), or generate unwanted genetic or pharmacological interactions (Gozzi et al., 2008) that can confound the mechanistic interpretation of rsfMRI signals. More importantly, the lack of well-characterized rsfMRI datasets in awake mice prevents a full understanding of how anesthesia-induced loss of consciousness affects the functional architecture of rsfMRI networks with respect to awake conditions in this species. This area of research is of especial significance, given the increasing interest in the study of how the dynamic structure of rsfMRI activity evolves and reconfigures as a function of brain state.

Influential investigations in wakeful and anesthetized primates and humans have uncovered hallmark dynamic features that are strongly biased towards high and low levels of consciousness, respectively. Specifically, loss of consciousness has been associated with a partial reorganization of long-range functional connectivity, disappearance of anticorrelated cortical states and a repertoire of dynamic states dominated by rigid functional configurations tied to the underlying anatomical connectivity (Barttfeld et al., 2015; Demertzi et al., 2019). By contrast, conscious wakefulness has been associated with greater global integration and inter-areal cross-talk (Demertzi et al., 2019), anticorrelations between the activity of different brain regions (Barttfeld et al., 2015; Huang et al., 2020), and a more flexible repertoire of functional brain configurations departing from anatomical constraints (Barttfeld et al., 2015; Demertzi et al., 2019). Interestingly, some of these network features have been proposed to constitute a putative “signature of consciousness” that exhibits significant evolutionary conservation in primates and humans (Barttfeld et al., 2015; Demertzi et al., 2019). However, it remains unclear whether and how analogous dynamic rules govern the spatiotemporal organization of spontaneous fMRI activity in the awake mouse.

Leveraging a robust protocol for rsfMRI in awake head-fixed mice, here we provide a fine-grained description of the functional topography and dynamic structure of rsfMRI networks in wakeful animals. By comparing awake network features with those obtained in anesthetized states, we find that rsfMRI networks in wakeful mice are topologically configured to maximize integration between neural communities, and exhibit topographically unique co-activation patterns encompassing a dominant involvement of arousal-related nuclei, anti-coordination between visual-auditory and default-mode-network areas, and idiosyncratic temporal transitions towards distinctive network attractor states. These results reveal that rsfMRI activity in awake mice is critically shaped by state-specific involvement of basal-forebrain arousal systems, and exhibits a stereotypic spatiotemporal structure that reconstitutes foundational principles of conscious network states in higher mammalian species.

## Results

### Implementation of resting state fMRI in awake head-fixed mice

Prompted by the emerging use of head-fixation to reduce motion-related artefacts in rodent neuroscience studies (Guo et al., 2014), we devised a simple protocol for awake mouse rsfMRI via the surgical implantation of plastic headposts, followed by gentle and progressive restraint onto a custom-made cradle (Fig. 1). After recovery from surgery, we subjected mice to progressive operator handling and mock scanning sessions of increasing length with the aim to acclimate them to the procedure and minimize restraint-induced stress (Fig. 1). Mice undergoing the habituation procedure did not experience substantial weight loss (Fig. S1), and exhibited a growth rate only marginally lower than that observed in co-housed control littermates not subjected to any experimental procedure (2-way ANOVA, group p = 0.39, session x group p = 0.15). We also measured plasma corticosterone levels in surgically implanted animals during the initial acclimation period (handling stage) and immediately after the first MRI scan at the end of habituation period. We next compared the resulting levels with those obtained in co-housed littermates under baseline conditions (no experimental procedure), or after a routine behavioral assessment (open field test, Fig. S1D-E). We found statistically higher level of corticosterone in the fMRI awake group under both conditions (baseline, and rsfMRI or OFT, 2-way ANOVA, group p = 0.02, session p < 0.001), however no group X session interaction (p = 0.90), suggesting that the acclimation and scanning procedure is overall only marginally more stressful than routine behavioral testing. Moreover, a comparison of measure corticosterone levels with those previously reported by others under different conditions or manipulations (Fig. S1E) revealed that the corticosterone levels associated with awake rsfMRI scanning were ca. 2-5 fold lower than those elicited by unhabituated acute immobilization, 2-fold lower than anesthesia-stimulated corticosterone release, and broadly comparable to the amounts elicited by natural circadian excursion, or induced by similar habituation protocols employed for awake MRI imaging in the mouse (Fig. S1E). Importantly, mice who underwent an open field test after fMRI scanning did not reveal any stress-related phenotype when compared to co-housed littermates not subjected to any restraint or habituation procedures (Fig. S1F, p > 0.31, all metrics). The results of these investigations suggest that the acclimation procedure is well-tolerated, and that stress-response produced by our habituation and awake fMRI scanning procedure is marginal.

**Figure 1.**
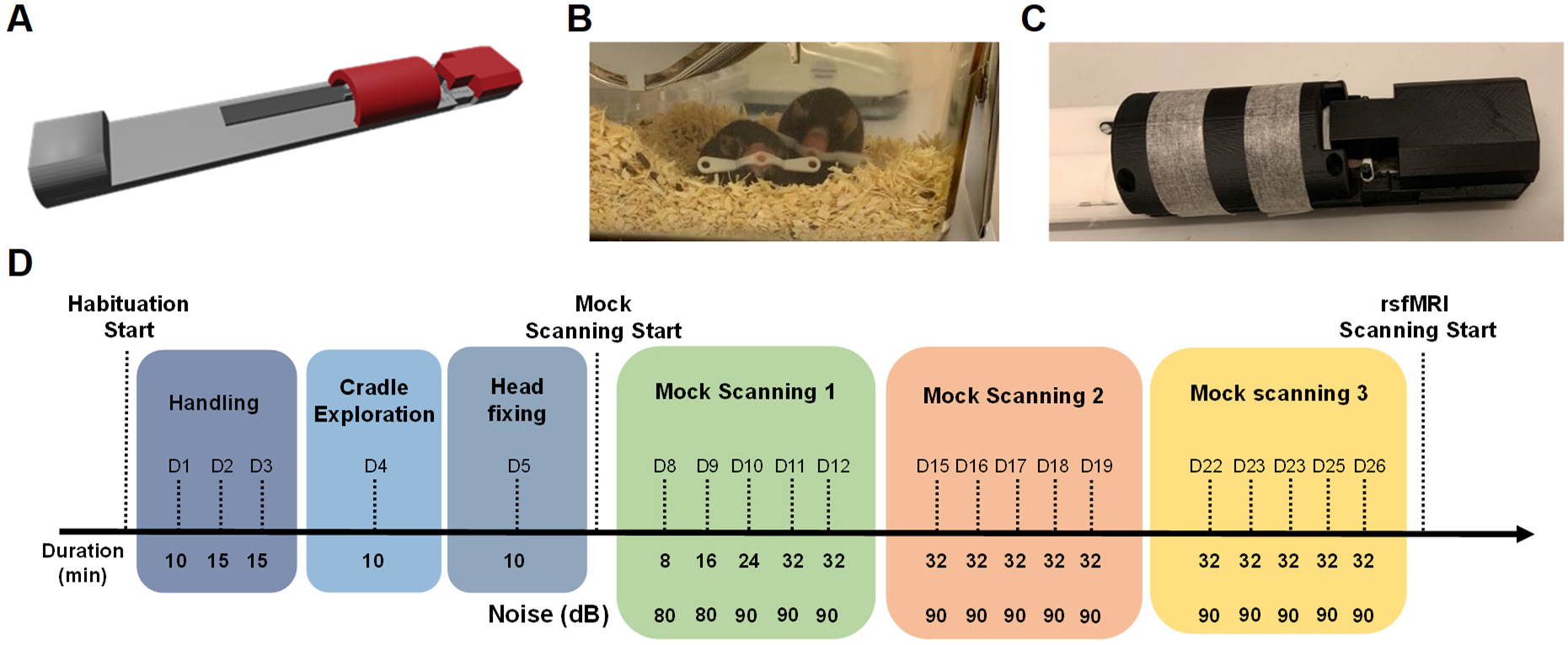
Acclimation protocol for awake rsfMRI. A) Three-dimensional render of the custom mouse cradle used for rsfMRI acquisitions (Ugo Basile S.r.L). B) Plastic headpost used for head fixing. C) Restraint apparatus upon and coil positioning. D) Experimental timeline for habituation protocol. D1 habituation started 10 days after head-post surgery.

### rsfMRI networks in awake mice exhibit focal functional reconfiguration

Using this protocol, we acquired 32-min long rsfMRI timeseries in n = 10 awake C57Bl6/J mice. To map the functional organization of rsfMRI networks in awake conditions, we systematically probed rsfMRI connectivity networks via seed-based correlation mapping. This analysis revealed robust interhemispheric and antero-posterior rsfMRI synchronization (i.e. “functional connectivity”), including the presence of distributed networks anatomically recapitulating rsfMRI system previously described in lightly anesthetized mice (Sforazzini et al., 2014; Grandjean et al., 2020; Whitesell et al., 2021). The identified systems include a default mode network (DMN); a salience (insular-cingulate) network; a sensory-motor latero-cortical network (LCN); a visual-auditory latero-posterior network (LPN), plus a number of subcortical sub-systems, including dorsal (caudate-putamen) and ventrostriatal networks (mesolimbic pathway); a dorsal hippocampal network and a widely-distributed olfactory and amygdaloid network (Fig. 2).

**Figure 2.**
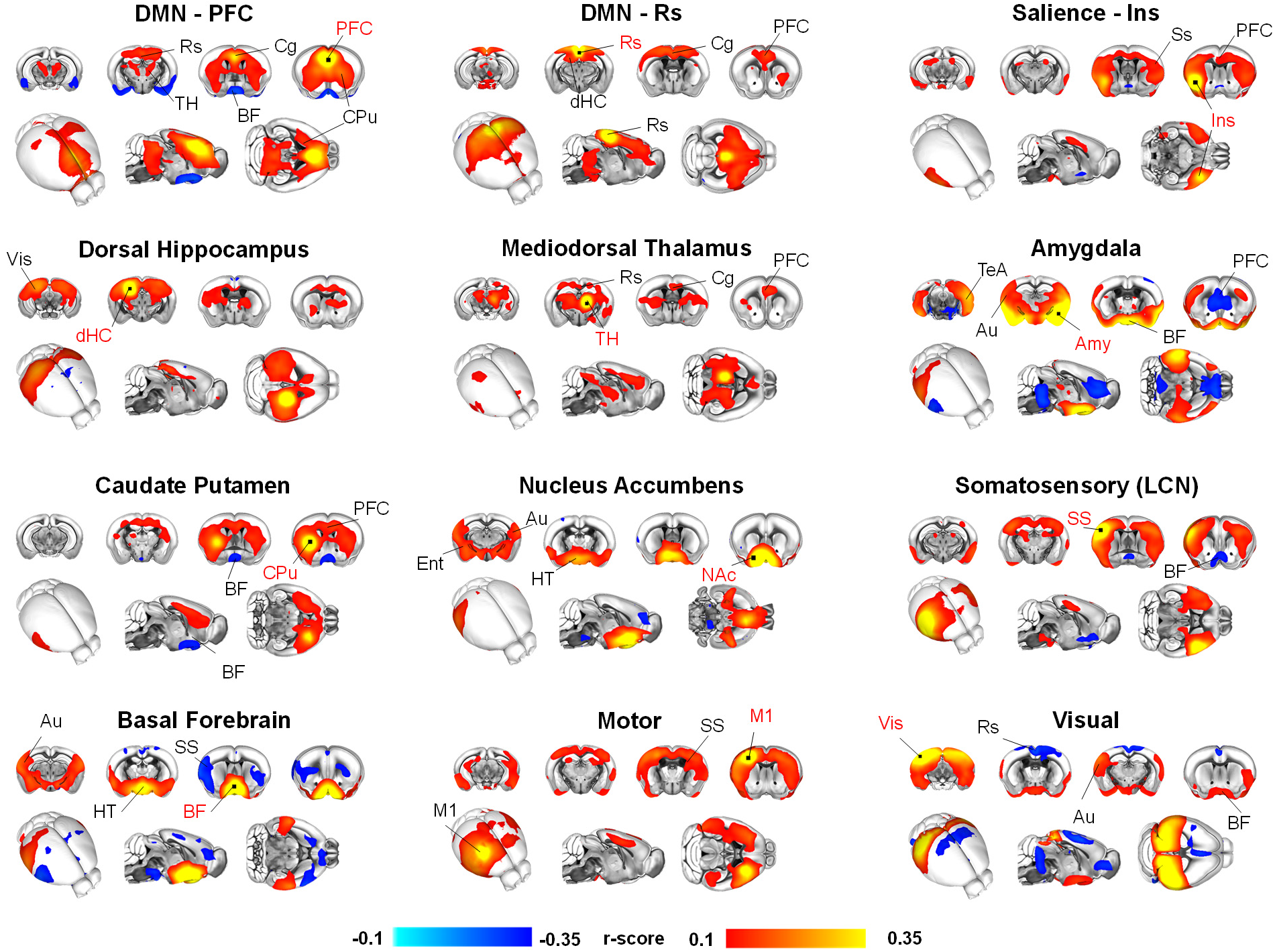
rsfMRI connectivity networks in the awake mouse brain. Each panel represents the averaged seed-based correlation map across n = 10 awake subjects, thresholded to voxels with significant connectivity (two-tailed T-test, p < 0.01, FWER cluster corrected with defining threshold T = 2.8). Seed location is indicated by red lettering. [DMN: Default mode network, PFC: Prefrontal cortex, Cg: Cingulate cortex, Rs: Retrosplenial cortex, TH: thalamus, CPu: Caudate putamen, Ins: Insula, dHC: dorsal Hippocampus, Ent: Entorhinal cortex, Au: Auditory cortex, M1: Motor Cortex, SS: Somatosensory cortex, V1: Visual, BF: Basal forebrain, Amy; Amygdala, NAc: Nucleus Accumbens, HT: Hypothalamus]

While the functional organization of rsfMRI networks in awake mice appears to broadly reconstitute key architectural features previously mapped in anesthetized mice, focal or more subtle state-dependent differences in the topography and organization of rsfMRI network activity may uncover dynamic features and functional substrates representative (and predictive of) wakeful, conscious states in lower mammalian species. To investigate this aspect, we spatially compared rsfMRI network topography in our awake scans with those previously acquired in a separate cohort of age-matched anesthetized C57Bl6/J mice (Gutierrez-Barragan et al., 2019). Interestingly, this comparison revealed a set of focal statedependent differences in the extension and anatomical organization of rsfMRI connectivity networks (Fig. 2AB). First, we found that rsfMRI networks in awake animals exhibited robust functional anti-coordination between some of the probed regions and their long-range substrates. Anti-coordination was especially prominent between medial prefrontal cortex (PFC) and olfactory regions, as well as between visual-auditory areas and midline regions of the DMN. The observed anti-correlation was accompanied by a reduced spatial extension of the DMN in awake mice, where a clear segregation of midline corticolimbic and visuo-auditory postero-lateral portions of the DMN was apparent. Moreover, in awake mice ventral forebrain area (e.g. diagonal band, hypothalamus, nucleus accumbens) were part of an extended highly-synchronous network that exhibited only marginal functional coupling in anesthetized subjects. More nuanced network-specific differences in rsfMRI connectivity strength were also apparent, with evidence of reduced cortico-cortical coupling in the DMN and LCN, and stronger connectivity in visual-auditory and basal forebrain areas in awake versus anesthetized animals.

These results were corroborated by whole-brain voxelwise mapping of interareal connectivity via correlation matrices (Fig. 3C-D). To help anatomical interpretability of this analysis, we organized voxels into axonal connectivity modules recently identified in the mouse brain (Fig. S2, Coletta et al., 2020). The resulting fine-grained partitioning (Fig. 3D) confirmed that in awake animals (a) the DMN reconfigures into two segregated submodular division (midline and sensory-PLN); (b) connectivity within basal forebrain systems, where most ascending neuromodulatory systems are located, is dramatically increased; (c) rsfMRI network activity exhibits higher functional anti-coordination (Fig. S2B), uncovering relations between networks that are not present in anesthetized conditions, such as the inverse coupling between midline and PLN components of the DMN (Fig. 3C-D). Corroborating these findings, an extension of our comparisons to a third cohort of mice anesthetized with a different anesthetic regimen (e.g. isoflurane plus medetomidine, Grandjean et al., 2020) replicated these topographical features (Fig. S2E-G), suggesting that they are not the result of a specific pharmacological mechanism, but can more broadly reflect state-dependent reconfiguration following anesthesia-induced loss of responsiveness. Taken together, these investigations document that wakeful states in the mouse lead to a focal rsfMRI network reconfiguration dominated by increased basal forebrain-cortical coupling, and anti-coordination between sensory and latero-posterior DMN associative cortices.

**Figure 3.**
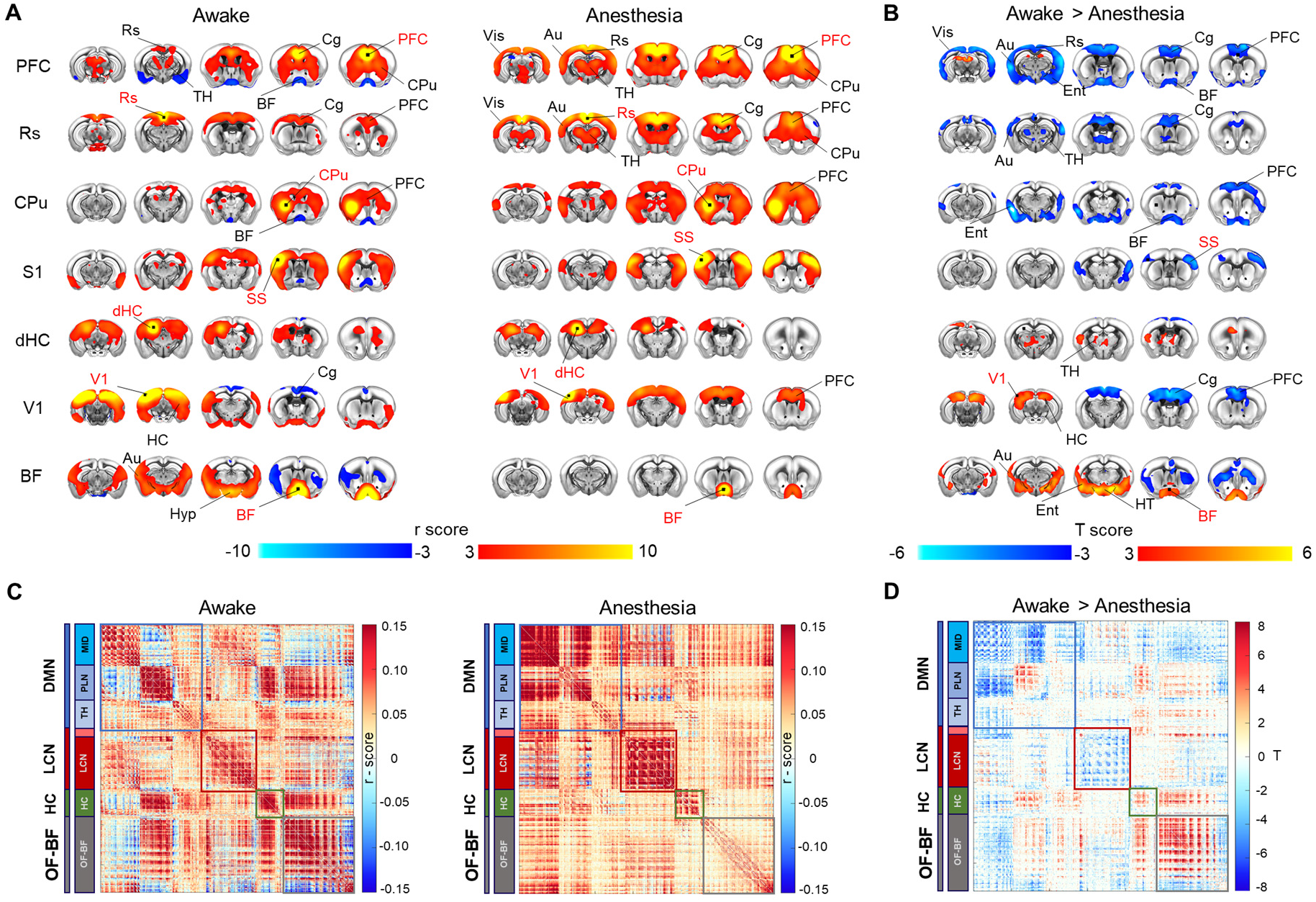
rsfMRI network topography in the awake and anesthetized mouse brain. **A)** Group averaged seed-based correlation maps in awake (N = 10, left) and halothane anesthetized (N = 19, right) mice, thresholded to voxels with significant connectivity (one sample T-test, p < 0.01, cluster corrected with defining threshold T = 2.8). **B)** Between group connectivity differences (two-sample T-test, p < 0.01, FWER corrected with defining threshold T=2.8). **C)** Group averaged voxel-wise FC matrices for each condition, with voxels organized within axonal connectivity modules (Coletta et al, 2020), and further partitions into network submodules. The sub-module corresponding to thalamic voxels within the LCN is not labeled (light-red block). **D)** Significant between condition FC differences at the voxel level (two-tailed two-sample T-test). [DMN: Default mode network, LCN: Latero-cortical network, HC: Hippocampal network, OF-BF: olfactory-Basal forebrain network, PFC: Prefrontal cortex, Cg: Cingulate cortex, Rs: Retrosplenial cortex, TH: thalamus, CPu: Caudate putamen, Ins: Insula, dHC: dorsal Hippocampus, Ent: Entorhinal cortex, Au: Auditory cortex, M1: Primary Motor, SS: Somatosensory cortex, V1: Visual, BF: Basal forebrain, Amy; Amygdala, NAc: Nucleus Accumbens, HT: Hypothalamus]

### rsfMRI connectivity in awake mice show increased between-network communication

The observation of areas of negative correlation in the awake mouse brain is of great interest, as similar findings have been suggested to serve as a putative signature of fMRI network activity in conscious states in other mammalian species such as marmosets (Hori et al., 2020a), macaques (Barttfeld et al., 2015) and human (Demertzi et al., 2019; Esfahlani et al., 2020). Importantly, these reports also showed that network configuration in anesthetized and wakeful animals may be characterized by different anatomical organization, with unconscious state being more tied to the anatomical map, and awake brain networks exhibiting topographical departure from their underlying anatomical architecture. To investigate whether similar principles would apply to the mouse brain, we used a graph theoretical approach to probe the relationship between structural and functional connectome in wakeful and anesthetized animals. To this aim, we leveraged a recent anatomical partition of the voxelwise mouse axonal connectome into four macro-communities that spatially reconstitute macroscopic network systems of the mouse brain ((i.e. the DMN, the LCN, the hippocampus and olfactory-basal forebrain areas, Coletta et al., 2020). A graphic representation of the functional connectome with respect to these axonal communities (Fig. 4A) revealed that, departing from the modular partitioning of the axonal connectome, fMRI networks in awake subjects exhibit greater inter-areal communication than corresponding anesthetized state. In keeping with this notion, structure-function correspondence was significantly lower in awake animals compared to anesthetized subjects (p<0.01, Mann-Whitney test, Fig. 4B). Formal quantifications of rsfMRI network connectivity strength corroborated these qualitative observations (Fig. 4C), revealing dramatically increased between-network connectivity in awake mice, a finding that was especially prominent between basal forebrain and cortico-hippocampal areas (p< 0.01, Mann-Whitney test, FDR corrected). Importantly, analogous features were observed when we contrasted awake rsfMRI data with those obtained in isoflurane-medetomidine anesthetized animals (Fig. S2, H-J), hence supporting a possible generalization of this finding to other anesthetic regimens. These findings recapitulate prior observations in conscious primates (Barttfeld et al., 2015), suggesting that, departing from the underlying structure of the axonal connectome, in the awake mouse brain rsfMRI network activity topologically reconfigures to maximize cross-talk between cortical and subcortical neural systems.

**Figure 4.**
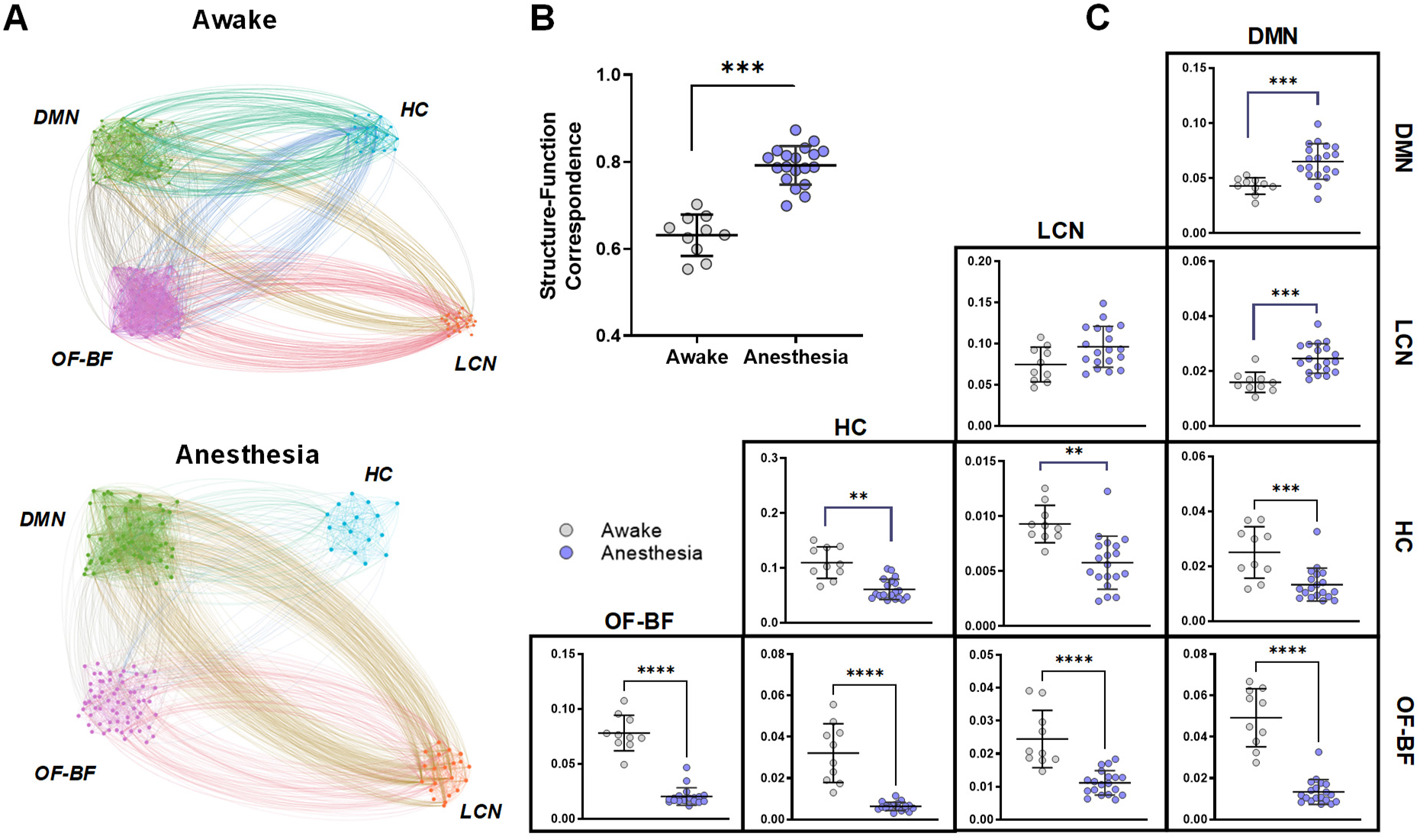
Structure-function relationship in awake and anesthetized states. **A)** Graphic representation of rsfMRI connectivity within and between previously described axonal modules of the mouse brain (DMN, LCN, HC, OF-BF, from Coletta et al., 2020). Each cluster of nodes represents a subset of anatomically-defined ROIs within the corresponding module. Nodes have been empirically arranged to maximize figure legibility. B) Structure-function correspondence in awake (N = 10) and anesthetized mice (N = 19). Between-group differences were assessed with a Mann-Whitney test (p < 0.05). C) Quantification of within (diagonal) and between (off-diagonal) network functional connectivity (*p<0.05, ** p<0.01, *** p<0.001, **** p<0.0001, Mann-Whitney test, FDR corrected). [DMN: Default mode network, HC: Hippocampus, OF-BF: Olfactory and basal forebrain, LCN: latero-cortical network].

### rsfMRI dynamics in awake mice exhibits unique coactivation topography

Our investigations revealed focal functional reconfiguration in “static” (i.e. time-averaged) rsfMRI networks in awake conditions. Prompted by the identification of dynamic connectivity signatures of consciousness in primates and human (Barttfeld et al., 2015; Demertzi et al., 2019; Huang et al., 2020), we hypothesized that the observed static network changes could similarly reflect state-dependent differences in the underlying dynamic structure of rsfMRI. To this aim, we decomposed rsfMRI activity into six dominant recurring co-activation patterns (CAPs), as these have been recently shown to govern rsfMRI dynamics in the mouse brain (Gutierrez-Barragan et al., 2019), and their dynamics has been found to be predictive of conscious states in humans (Huang et al., 2020).

In keeping with previous investigations (Gutierrez-Barragan et al., 2019), all the highlighted CAP topographies encompassed recognizable network systems of the mouse brain (Fig. 5A, and opposing spatial configurations, delineating three dominant CAP-anti-CAP pairs (CAP 1-2; 3-4 and 5-6, Fig. 5B) capturing peaks and throughs of fluctuating BOLD activity. Interestingly, while the overall anatomical organization of CAPs appeared to be comparable in awake and anesthetized conditions, distinctive statedependent topographical differences in the anatomical organization of the corresponding BOLD coactivation patterns were apparent (Fig. 5B). An especially notable finding was the observation of robust coactivation of arousal-related basal forebrain nuclei in CAPs 5-6 of awake animals. This spatial signature was associated with a weaker but extended coactivation of thalamic and latero-posterior visuo-auditory regions, which exhibited anticoordinated peaks of BOLD activity in DMN and visual areas in awake state. CAPs 3 and 4 in awake mice similarly showed enhanced fluctuations in basal forebrain areas, together with the diffuse involvement of striatal and thalamic substrates. Finally, CAPs 1 and 2 in awake mice were characterized by focally dampened peaks of BOLD activity in midline areas of the DMN and in the LCN. These topographic differences show that rsfMRI dynamics in the awake state is characterized by a distinctive coactivation of arousal-related subcortical nuclei and additional network components (e.g. PLN, midline DMN) that we found to be differentially configured in the static connectome of awake and anesthetized animals (cf. Fig 2). This finding corroborates a tight link between CAP dynamics and the ensuing time-averaged network activity (Gutierrez-Barragan et al., 2019), suggesting that the functional architecture of the “static” rsfMRI connectome is critically shaped by its underlying coactivation structure.

**Figure 5.**
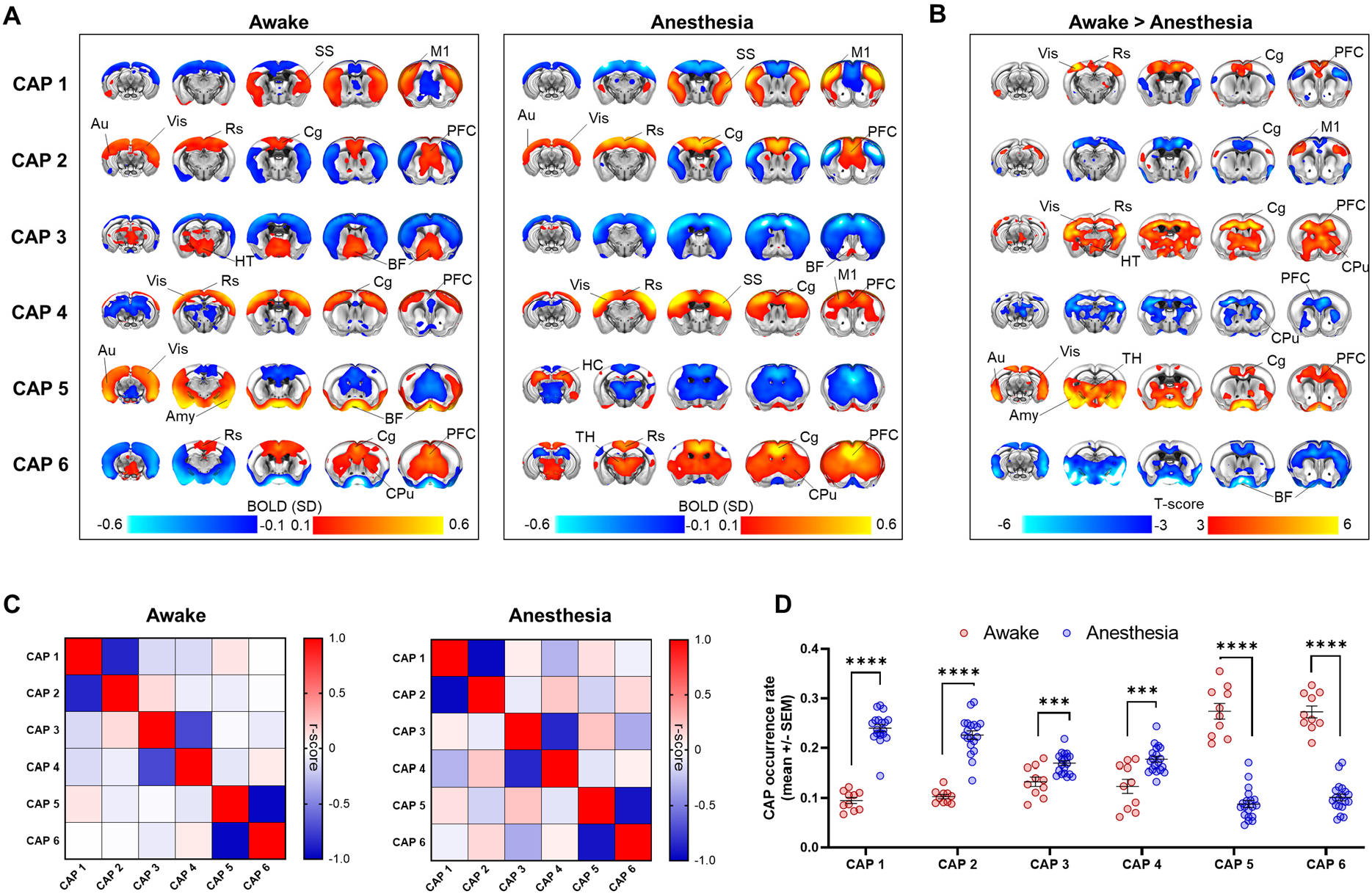
Coactivation pattern (CAP) topography and occurrence in the awake and anesthetized mouse brain. **A)** CAP topography in awake (N = 10, left) and anesthetized mice (halothane, N = 19, middle; T-test, p < 0.01, FWER cluster corrected with defining threshold T=2.8). **B)** Corresponding awake > anesthesia difference maps (T-test, p < 0.01, FWER cluster corrected with defining threshold T=2.8). **C)** Between CAP spatial similarity (Pearson’s correlation) in awake and anesthetized mice. Note the presence of clear CAP anti-CAP pairs. D) Quantification of CAP occurrence rates in awake and anesthetized mice (***p< 0.001, ****p< 0.0001, T-test, FDR corrected). [PFC: Prefrontal cortex, Cg: Cingulate cortex, Rs: Retrosplenial cortex, TH: thalamus, CPu: Caudate putamen, Ins: Insula, HC: Hippocampus,, Au: Auditory cortex, M1: Primary Motor, SS: Somatosensory Cortex, V1: Visual, BF: Basal forebrain, Amy; Amygdala, NAc: Nucleus Accumbens, HT: Hypothalamus]

Further supporting this notion, it has been recently shown that most variance in static rsfMRI connectivity is explained by a small fraction of fMRI frames exhibiting the highest co-fluctuation amplitude (Esfahlani et al., 2020). This observation suggests that CAPs might similarly represent peaks (and troughs) of BOLD activity that critically shape the structure of the static rsfMRI connectome. To test this hypothesis, we generated CAP cofluctuation matrices for both awake and anesthetized conditions (Fig. S3) and compared their mean topography with that of the corresponding time-averaged, “static” functional connectivity. This comparison yielded a spatial correlation r = 0.79 (R^2^ = 0.62) and r = 0.78 (R^2^ = 0.61) for awake and anesthetized rsfMRI timeseries, respectively. Similarly high spatial correlation was found when the approach was applied to medetomidine-isoflurane anesthetized animals (r = 0.69, R^2^ = 0.49). These results expand recent investigations (Esfahlani et al., 2020), by showing that the peaks of BOLD activity captured by CAPs account for a dominant fraction of variance in time-averaged static rsfMRI connectivity. They also suggest that CAPs, and their unique state-dependent functional configuration, crucially shape the topography and dynamics of rsfMRI networks mapped in awake and anesthesia-induced loss of responsiveness.

### Prevalence of coactivation patterns distinguishes awake and anesthetized states

Human and primate research has shown that conscious states are associated with rsfMRI connectivity signatures characterized by the dominant occurrence of stereotypical functional configurations (Barttfeld et al., 2015; Demertzi et al., 2019; Huang et al., 2020). These findings led us to investigate whether analogous changes in CAP dynamics could underlie the network reconfiguration observed in wakeful mice. To this aim, we first probed CAP oscillatory dynamics by quantifying the fractional amplitude of low-frequency fluctuations (fALFF, Fig. S4C) in the spectral power of each CAP’s timeseries (Fig. S4A-B). This analysis revealed that, although both wakeful and anesthetized states exhibit broadly comparable infra-slow [0.01 −0.03 Hz] dynamics, the power of these slow fluctuations was significantly higher under anesthesia (p< 0.05, Mann-Whitney test, all CAPs except CAP4). We next computed the distribution of GS-phases at the occurrence of each CAP within infraslow (0.01-0.03Hz) fMRI GS cycles, as previous investigations revealed that in anesthetized conditions CAP occurrence is phase locked to GS cycles (Gutierrez-Barragan et al., 2019). Interestingly, we found that CAP occurrence in awake states is also locked to GS infraslow cycles (Rayleigh test, p < 0.001, FDR corrected, Fig. S4D). However, CAP3 and 4 did not exhibit opposite phase occurrence as seen in anesthetized animals, but showed instead concordant preferred occurrence in antiphase with that of GS cycles. Moreover, phase occurrence of awake CAPs 5 and 6 was in antiphase with that of the corresponding CAPs in anesthetized animals. These results document that, while CAPs in both anesthetized and awake conditions occur at specific phases of GS fluctuations, significant state-dependent changes in the preferred phase occurrence of these coactivation patterns exist.

Lastly, to assess whether any of the observed CAPs could be predominantly associated with a specific brain state, we compared CAP occurrence in awake mice and anesthetized animals (Fig. 5C). This analysis revealed dramatic state-dependent differences in CAP prevalence. Specifically, occurrence of CAPs 1 and 2 (and to a much lower extent, CAPs 3 and 4) was significantly higher (ca. 2-5 fold) in anesthesia than awake states. Conversely, CAPs 5 and 6 were prominently more frequent (ca. 3-fold) in the awake state (p<0.001, two-tailed, two-sample T-test, FDR corrected). These result show that in the mouse prevalence of CAPs is critically different in awake and anesthetized states. Importantly, very similar CAP occurrences were observed in animals anesthetized with isoflurane-medetomidine (Fig. S5), supporting a possible generalization of this observation across anesthetic conditions. It should also be noted that the CAPs that are prevalent in the awake state (CAP5 and 6) exhibit distinctive anti-coordinated engagement of visual-auditory regions (PLN) and DMN areas (Fig. 5A and S3A). This finding recapitulates similar patterns of regional anticoordination in higher mammalian species (Barttfeld et al., 2015; Demertzi et al., 2019), pointing at a putative evolutionarily-conserved network signature predictive of wakeful, conscious states in the mammalian brain.

### rsfMRI networks in awake mice exhibit unique temporal structure

Prompted by recent evidence of stereotypic CAP transition trajectories in conscious humans (Huang et al., 2020), we next investigated whether similarly different state-dependent temporal trajectories could be identified in awake and anesthetized mice. To unravel sequential transitions within the awake and anesthetized states, we modelled CAP timeseries as a Markov process including both the transition probability between CAPs as well as self-transitions (e.g. persistence probability). Consistent with the slow dynamics of CAPs, we found that all persistence probabilities in both awake and anesthetized conditions were highly significant when compared to a null-hypothesis distribution with randomly permuted CAP sequences (p < 0.001, all CAPs). In agreement with CAP occurrence results, between-state comparisons next showed that CAPs 1-2 and 3-4 exhibit higher persistence in the anesthetized state, while CAPs 5-6 show increased persistence in awake conditions (Fig. 6A-C, top row).

**Figure 6.**
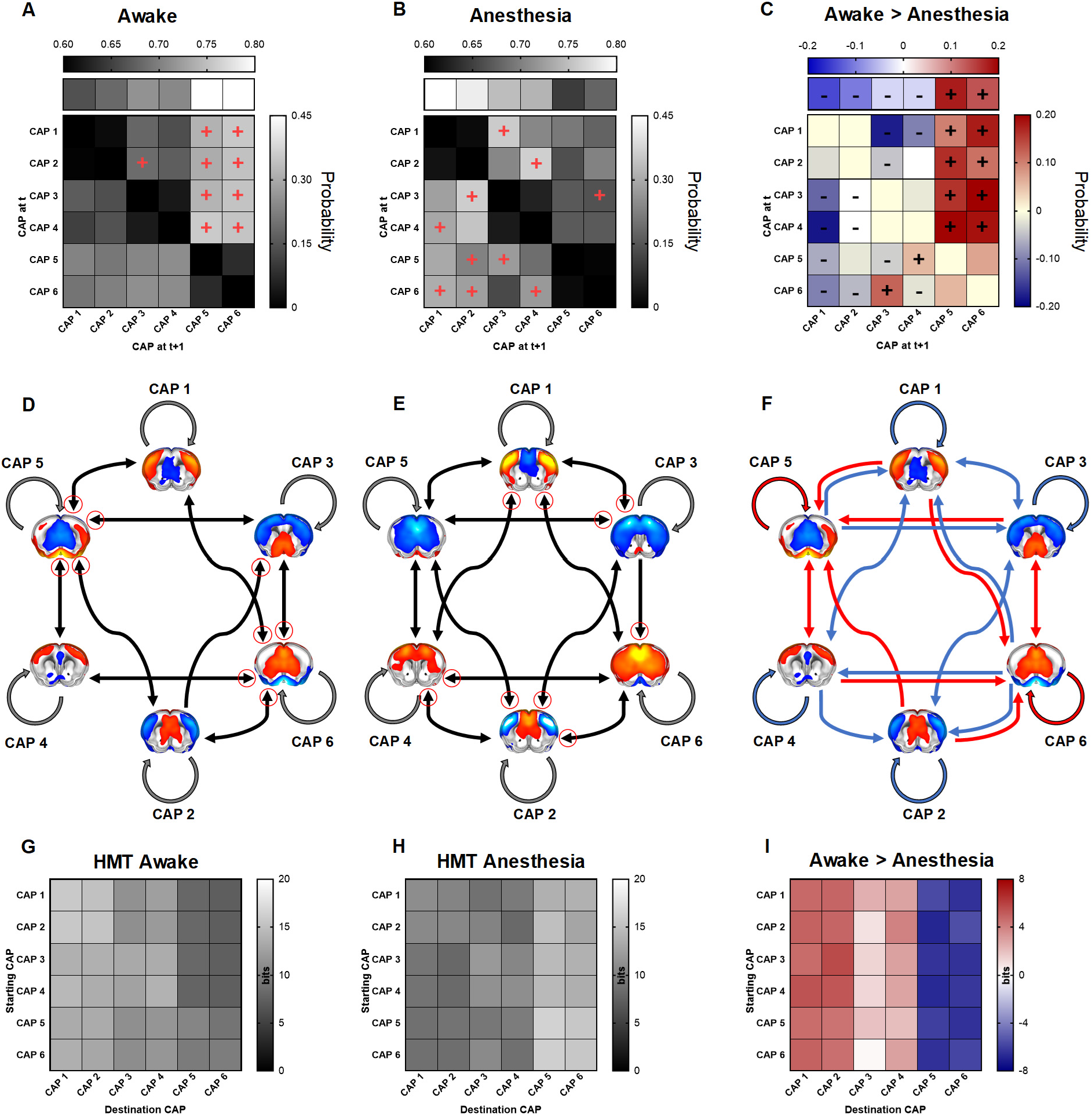
CAP transition probabilities and corresponding temporal trajectories. **A-B)** Persistence (top row) and transition (off-diagonal) probability in awake (A, N = 10) and anesthetized (B, N = 19) mice. Significant preferred transitions *P_ij_* >*P_ji_* are denoted with a red “+” sign. Directional transitions are depicted if the incoming transition *P_ij_* is significant. **C)** Transition probability differences matrix. Statistically significant between-group differences are denoted with a “+” sign for transitions higher in awake, and conversely with a “−” sign, if higher in anesthesia. **D-E)** Graph representation of significant persistence (round self-connecting arrows) and transition (black arrow heads) probability for awake and anesthesia conditions, respectively. Significant transitions (p<0.001, 1000 surrogate timeseries) are depicted with an arrowhead, while directionally dominant transitions are denoted with a red circle. **F)** Significant persistence and transition probability differences between groups. Red arrows indicate higher transition probabilities in awake mice, and blue arrows in anesthetized mice. **G-H)** Entropy of Markov trajectories (HMT) in awake (G) and anesthetized mice (H). Higher entropy of a trajectory indicates lower accessibility of a CAP destination from a starting CAP. **I)** Between-group differences in HMT. Positive elements represent CAP trajectories with lower accessibility in the awake state as compared with anesthesia, while negative elements are trajectories with facilitated access in awake. Only paths from CAPs 2 and 6 to CAP 3 did not reach statistical significance at p<0.001 (1000 surrogate timeseries).

To portray the temporal trajectory of CAP transitions in awake and anesthetized mice, we next computed, and graphically depicted, the corresponding transition probability matrices (Fig. 6A-F). These investigations revealed stereotypic CAP trajectories in awake and anesthetized conditions. Specifically, we found that in awake mice CAPs 5 and 6, besides being the most recurrent (Fig. 5C) and persistent CAPs, were also configured as dominant attractors state, i.e. they are the most probable destination from all other CAPs. Conversely, temporal transitions under anesthesia where more distributed, with both pan-cortical CAPs 3-4 and DMN-LCN CAPs 1-2 (i.e. the two CAP pairs with the highest occurrence in this brain state), emerging as the most prominent CAP attractors (Fig. 5C). Importantly, very similar temporal signatures were observed in mice anaesthetized with medetomidine-isoflurane (Fig. S6A-C), corroborating the robustness of this finding. Moreover, our results were confirmed after performing 500 split-half resampling of the sequences (r > 0.96; Fig. S6G) and also after comparing the group concatenated probabilities to each single mouse (r > 0.85 all mice, with one exception r = 0.69; Fig. S6H). These additional analyses strongly corroborate the robustness and state-specific nature of the identified temporal transitions signatures identified.

To investigate whether the observed temporal transitions could be associated with statedependent differences in the complexity of trajectories between each pair of CAPs, we computed the entropy of Markov trajectories (HMT) for each transition probability matrix. This metric provides an estimate of the “accessibility” of each CAP from another one: low descriptive complexity (i.e. entropy close to 0) from a starting point (initial CAP) to its destination (final CAP) indicates an almost deterministic direct path from CAP i to CAP j (high accessibility). In contrast, high entropy values (i.e. close to 1 bit) imply a higher uncertainty, and lower accessibility, as the trajectory encompasses random steps through different CAPs to reach its destination. Consistent with our CAP occurrence results and the temporal structures described above, out investigations revealed that CAPs 5-6 are the two temporal states with the highest accessibility in awake animals (Fig. 6G-H), and conversely those exhibiting the lowest accessibility in anesthetized states. CAPs 1-2 and 3-4 required instead more elaborate transition trajectories in awake conditions, but had higher accessibility in anesthetized animals (Fig. 6I). Supporting the generalizability of our findings to other anesthetic regimens, a remarkably similar CAP transition signature was observed in isoflurane-medetomidine anesthetized mice (Fig. S6D-F). Moreover, our results were robust after repeating the calculations both with 500 split-half resampling iterations of each dataset (Fig. S6I), and at the single-mouse level (Fig. S6J). Taken together, these results indicate that rsfMRI activity in the awake mouse exhibits stereotypic temporal trajectories and increased accessibility to a state-dominant network configuration (CAPs 5 and 6) characterized by a critical engagement of arousal-related basal forebrain areas, and anti-coordination between DMN and visual regions.

## Discussion

Leveraging a robust protocol for rsfMRI mapping in awake mice, we investigated the functional architecture and dynamic organization of rsfMRI activity in wakeful and lightly anesthetized mice, with the aim of determining how the ensuing network architecture reconfigures as function of the underlying brain state. Our investigations revealed that rsfMRI activity in awake mice undergoes focal topographic reconfiguration entailing the presence of network-specific regional anticorrelation, heightened connectivity in arousal-related basal forebrain areas and greatly increased between-network cross-talk, departing from the rigid structure of the axonal connectome. Notably, these changes were associated with remarkably distinct dynamic patterns of signal coordination as assessed with a deconstruction of rsfMRI activity into recurring coactivation patterns. Specifically, we found that rsfMRI activity in wakeful animals exhibits a largely dominant occurrence of CAPs encompassing arousal-related forebrain nuclei, as well as features recently described to be predictive of consciousness in higher mammalian species, such as transient anticorrelation between visual-auditory and DMN areas (Barttfeld et al., 2015; Demertzi et al., 2019), and a stereotypic temporal transitions driven by a set of more accessible and persistent temporal states (Huang et al., 2020). These results suggest that rsfMRI activity in awake mouse is critically shaped by a state-specific involvement of basal-forebrain arousal systems, and that its dynamic structure recapitulates evolutionarily-relevant principles described in higher mammalian species.

Adding to recent attempts to map rsfMRI networks in the mouse (Yoshida et al., 2016; Tsurugizawa and Yoshimaru, 2021), our work describes a protocol enabling the reliable implementation of rsfMRI imaging in awake head-fixed animals and provides a fine-grained description of the network structure in non-anesthetized rodents. Our investigations show that the overall static architecture of rsfMRI networks in awake mice reconstitutes organizational principles observed in anesthetized conditions, including the presence of distributed systems such as a DMN, a LCN and a salience-like network. Analogous between-state correspondences in anesthesia and awake conditions have been reported in rats (Liang et al., 2012a), primates (Xu et al., 2019; Hori et al., 2020b) and humans (Boveroux et al., 2010; Akeju et al., 2014), underscoring a tight relationship between the general spatial structure of spontaneous fMRI activity, and its underlying structural map (Alstott et al., 2009; Coletta et al., 2020).

The focal topographic reconfiguration observed in awake state is of interest as it highlights functional substrates that are critically sensitive to the effect of general anesthesia, and that as such may putatively govern a transition between stimulus-unresponsive to wakeful, conscious states in the rodent brain. In this respect, the observed heightened functional connectivity in basal forebrain-hypothalamic areas and their increased cross-talk with other cortical modules is fully consistent with the established role of these regions as key mediators of arousal and vigilance in the mammalian brain (Lee and Dan, 2012; Xu et al., 2015; Montani et al., 2020). As conscious perception relies upon the ability to integrate information across specialized communities of brain regions, our finding that rsfMRI network structure in awake states shows increased interareal cross-talk is important, as it reconstitutes in mice a dominant functional configuration that has been associated with conscious states in other species (Barttfeld et al., 2015; Ma et al., 2017). This finding is also consistent with the postulates of prevailing theories of consciousness (Dehaene and Changeux, 2011; Tononi et al., 2016), according to which functional networks that support awake, conscious states must exhibit global integration, evidenced in our data as greater functional coupling between cortical and sub-cortical network systems.

Our topographic investigations also corroborate the notion that during conscious wakefulness rsfMRI activity is distinctly characterized by the appearance of anticorrelation between the activity of different brain regions, a feature that is virtually absent in anesthetized conditions. It should be noted here that, because our rsfMRI timeseries were processed identically across all conditions and without regression of the global fMRI signal, this state-dependent change cannot be attributed to methodological artefacts (Murphy et al., 2009). Analogous observations have reported in rats (Liang et al., 2012b), primates (Barttfeld et al., 2015; Hori et al., 2020a) and humans (Demertzi et al., 2019), where they have been theoretically linked to the global neuronal workspace theory (Dehaene and Changeux, 2011). According to this view, different streams of information compete to engage widespread networks of regions via the mutual inhibition of activity at different cortical areas, leading to anticorrelated dynamics. Within this framework, the observed segregation of medial corticolimbic and postero-lateral visuo-auditory cortical portions of the DMN in awake states is of interest, as it highlights a focal, state-dependent network reconfiguration occurring in awake rodents (including rats, Liang et al., 2012b), that however does not appear to have a direct correlate in higher mammalians (Vincent et al., 2007), with the possible exception of new world primates (Liu et al., 2019). As this segregation affects a widely-distributed community of monosynaptic connections (Coletta et al., 2020; Whitesell et al., 2021), we speculate it could reflect a dominant configuration aimed to enable increased cortical information capacity (i.e. the ensuing number of discriminable activity patterns), in the otherwise poorly differentiated rodent posterolateral cortex (Alkire et al., 2008; Iurilli et al., 2012; Buckner and Krienen, 2013). Such a functional segregation of the posterior and midline DMN components may not be necessary in higher mammalian species, owing to a larger and more specialized cortical differentiation.

Our deconstruction of awake rsfMRI activity in recurring CAPs revealed a unique dynamic structure recapitulating foundational principles of network organization in conscious primates and humans, including (a) the identification of a largely state-dominant CAPs characterized by a rich topographical organization, not rigidly anchored to the axonal structure (Barttfeld et al., 2015) and by prominent anti-coordination between visual-auditory and DMN regions (Demertzi et al., 2019); (b) stereotypic temporal trajectories in which state-specific CAPs are configured as dominant network attractors (Huang et al., 2020). By contrast, anesthetized state was associated with dominant temporal states that are topographically shaped by the underlying axonal structure with extensively integrated DMN and visual-auditory regions (Gutierrez-Barragan et al., 2019; Coletta et al., 2020). These findings are consistent with prior theoretical conceptualizations of conscious network activity as a balanced recurrence of intermittent epochs of segregated and global brain synchronization (Demertzi et al., 2013). Above and beyond this, our results highlighted key state-dependent differences in the topographical organization of co-activation patterns, with evidence of distinctive involvement of basal forebrain, hypothalamic and thalamic areas in the awake, but not anesthetized, state. Owing to the tight link between CAP topography and the resulting patterns network activity (Gutierrez-Barragan et al., 2019), this finding is important as it suggests that rsfMRI network dynamics in awake states is critically shaped by basal forebrain and thalamic substrate areas, corroborating a previously postulated involvement of these regions to wakeful network organization (Alkire et al., 2008). Importantly, our investigations of CAP topography show that state-specific differences in rsfMRI dynamics may be accompanied but significant state-specific topographical reconfiguration of the ensuing coactivation patterns, leading to the identification of functional substrates that may critically shape network activity. The future extension of our analytic approach to map state-dependent topographic differences in higher species is warranted to investigate the predictive and evolutionarily validity of our observations across species and consciousness states.

A strength of our approach is the use of a translatable readout which allowed us to relate the observed state-specific changes to analogous investigation in higher mammalian species. However, some limitations need be recognized when extrapolating and comparing our findings across species and studies. First, the extent to which rsfMRI in habituated head-fixed mice (and conceivably primates - (Barttfeld et al., 2015) can recapitulate the quiet wakefulness state that characterize human rsfMRI studies remains unclear. Our corticosterone measurements are encouraging, as they show that stress response to our acclimation procedure was marginal. However wakeful imaging in head-fixed rodents may possibly entail arousal states that are not directly comparable to those attainable in humans. rsfMRI studies encompassing concurrent pupil-tracking (Pais-Roldán et al., 2020) may crucially help relate some of the functional signatures we describe here to arousal-state fluctuations. Second, the anesthetic regimens we used in our studies generally correspond to light anesthesia (Ferrari et al., 2012), and although they induce animal immobility and unresponsiveness, they might, at least conceivably, induce intermittent states of residual “consciousness” (Alkire et al., 2008). Similarly, the lack of internal state reporting in animals cautioned us to minimize use of “conscious state” when referring to the brain state associated with awake scanning. For this reason, all our references to conscious feature in the present work are the results from the indirect comparison of our findings with corresponding investigations in human studies. Notwithstanding these limitations, the degree of correspondence between the dynamic signatures we found in the mouse and those highlighted by corresponding primate and human research is remarkable, supporting a possible (yet cautious) extrapolation of these signatures across species.

In conclusion, we describe a robust protocol for rsfMRI mapping in awake mice, and report that the corresponding patterns of rsfMRI activity exhibit stereotypic spatiotemporal dynamics characterized by a dominant occurrence of coactivation patterns involving arousal-related nuclei, regional anticorrelation, and idiosyncratic temporal trajectories. These results suggest that dynamic structure of rsfMRI activity in the awake rodent brain recapitulates evolutionarily-relevant principles predictive of conscious states in higher mammalian species, and pave the way to the implementation of awake rsfMRI in this species.

## Acknowledgements

This study was funded by the European Research Council (ERC) under the European Union’s Horizon 2020 research and innovation program (#DISCONN; no. 802371 to A. Gozzi). A. Gozzi also acknowledges the support from the Brain and Behavior Foundation (NARSAD Independent Investigator Grant #25861), the NIH (1R21MH116473-01A1) and the Telethon foundation (GGP19177). The authors wish to thank all the members of the Gozzi lab for critically reading the manuscript.

## Materials and Methods

### Experimental Procedures

*In vivo* experiments were conducted in accordance with the Italian law (DL 26/214, EU 63/2010, Ministero della Sanità, Roma) and with the National Institute of Health recommendations for the care and use of laboratory animals. The animal research protocols for this study were reviewed and approved by the Italian Ministry of Health and the animal care committee of Istituto Italiano di Tecnologia (IIT). All surgeries were performed under anesthesia.

#### Animals

Adult (< 6 months old) male C57BL/6J mice were used throughout the study. Mice were group housed in a 12:12 hours light-dark cycle in individually ventilated cages with access to food and water ad libitum and with temperature maintained at 21 ± 1 °C and humidity at 60 ± 10%.

#### Experimental groups and datasets

A first group of mice (n=10, awake dataset) was subjected to the headpost surgery and habituation procedure and underwent awake fMRI image acquisitions as described below. The scans so obtained constitute the awake rsfMRI dataset we used throughout our study. Two additional groups of age matched male C57BL/6J mice were used as reference rsfMRI scans under anesthesia. The first group of animals (n=19, halothane dataset) was previously scanned in our laboratory under shallow halothane anesthesia, (0.75%, Gutierrez-Barragan et al., 2019). In the present work we have used this dataset as benchmark reference anesthesia dataset for all comparisons with awake fMRI networks, because the employed anesthesia regimen is well characterized (Sforazzini et al., 2016; Whitesell et al., 2021), it is representative of the network architecture observed with different anesthesia regimens in rodents (Grandjean et al., 2020), it exhibits rich spatiotemporal dynamics by preserving spectral properties of BOLD fluctuations (Gutierrez-Barragan et al., 2019) and it closely models the organization of the underlying axonal connectome (Coletta et al., 2020).

To probe a possible generalizability of our findings to other anesthetic conditions, we imaged a second, separate group (n=14) of mice under medetomidine-isoflurane anesthesia (0.05 mg/kg bolus and 0.1 mg/kg/h IV infusion, plus 0.5% isoflurane). While this anesthetic combination produces a non-physiological shift in the spectral components of BOLD fluctuation (Grandjean et al., 2014), it nonetheless represents the most-widely anesthetic mixture used in the mouse imaging community (Grandjean et al., 2020), and for this reason we used it a supplementary reference dataset. All the imaged mice were bred in the same vivarium and scanned with the same MRI scanner and imaging protocol employed for the awake scans (see below).

An additional cohort of n = 14 male mice was used to assess the effect of habituation and scanning on acute and chronic stress levels, as assessed with blood corticosterone level measurements, body-weight tracking, and an open field behavioral test. Within this cohort, n = 7 randomly selected “awake fMRI” mice were subjected to the same head-post surgery, habituation and awake scanning as our main rsfMRI dataset. Control littermates (n=7) were not implanted, habituated or fMRI scanned, but always left in their home cages with the exception of an initial in-cage blood corticosterone sampling (baseline, Fig. S1) and an additional one after an open field behavioral test. This group of mice was used to obtain reference baseline corticosterone levels timed to those obtained in the experimental group. The actual blood sampling timepoints corresponded to day 15 post-surgery in the fMRI awake cohort (handling, Fig. S1A), and day 39 post surgery, right after the first rsfMRI scan, which we carried out in the fMRI awake group only. Both set of mice were then subjected to the open field test to assess behavioral phenotypes indicative of chronic stress. Blood sampling was carried out at the end of the handling procedure session (fMRI awake), or in the home cage (control), and at the end of the MRI session (for awake mice) or open field test (for control mice). Housing was organized such that each cage contained a mix of both control and awake mice.

#### Headpost surgery

Head fixation during awake imaging was achieved using custom-made headposts (Ugo Basile, Italy; Figure 1B). A surgical procedure was performed to adhere the headposts to the skull. During surgery mice were anesthetized with isoflurane (5% induction and 2% surgery) and head fixed on a stereotaxic apparatus (Stoelting Co.) while body temperature was maintained with a heating pad at 37 °C. The fur on the head was removed and the skin was triple scrubbed with 70% ethanol. Once it was confirmed that the surgical anesthetic plane was maintained, a vertical incision (from a few mm anterior to bregma to the posterior end of the skull up to the eye level) was cut and the skin on top of the skull was removed. The headpost was cemented to the skull with a two-step procedure: first, dental cement (Superbond C&B kit, Dental leader) was applied to the skull surface; the headpost was pressed gently on top until the cement was partially dried (1-2 minutes). A second cement (Paladur, Kulzur s.r.l) was then applied to the skull and the headpost and allowed to dry for 5 minutes. Extra dental cement was applied around the edges to ensure complete coverage of all exposed bone. Suturing was not necessary as the skin naturally tightens around the craniotomy. Once the procedure was complete, mice were allowed to recover in their home cages with repeated monitoring. The headposts were well tolerated and showed no deterioration nor detachment over a 10-week examination window. After surgery, head post implanted mice could be group-housed without apparent direct damage to other headposts.

#### Habituation

A custom-made MRI-compatible animal cradle (Ugo Basile, Italy; Figure 1A, C) was designed such that the headposts could be secured to the cradle during scanning. A habituation protocol to acclimate the mice to immobilization and scanning was initiated 10-15 days after headpost surgery to allow sufficient time for recovery. The habituation timeline is depicted in Figure 1D, and was performed in two steps, entailing an initial stage of handling followed by extensive mock scanning. For the first 3 days (habituation day [D]1-3), 15 minutes daily were dedicated to acclimate the mice to the experimenter. On D4 and D5, mice were allowed to explore the cradle for 10 minutes, but on D5 the experimenter also would grab and hold the headpost by hand for a few seconds (during the 10 minutes) to acquaint the mouse with head fixation. Habituation to the mock scanner began on D8. During mock scanning, the headpost was secured to the cradle with plastic screws, a 3D printed copy of the RF-receive coil was placed over the head of the mouse, and the body was left unrestrained for the first two mock scanning habituation sessions. In the third mock scanning habituation session, the body of the mouse was gently restrained by taping its back to the cradle arc. Specifically, the mouse back was gently taped to the cradle in such as a way that the animal was able to breathe comfortably and make small movements below the solid edges of the cradle. Its tail was also taped to the base of the cradle but its forepaws and hindpaws were not. Cotton rolls were placed to reduce jaw and forepaw movement. A sound system played an audio recording of the EPI pulse sequence at increasing loudness to reach dB level recorded in the scanner bore. Figure 1D shows the duration and sound level for each daily session of mock scanning (D8-D27).

#### Open field test (OFT)

The open field test was adapted from existing protocols (Boulle et al 2014). Testing was carried out in a square arena (44 cmx 44 cm x44cm) with a homogenous 130 lux illumination subdivided in a central zone (24 cm width) and a peripheral zone (10 cm width). Mice were placed in the apparatus for a 20 minutes test period, during which total distance travelled (cm), immobile time (s), central distance (m) were automatically scored by Anymaze software (Stoelting Co). Behavioral data were analyzed performing an unpaired t-test between groups.

#### Plasma corticosterone measurement

The overall procedure of blood collection was no longer than 5 min and all blood collections (and corresponding tests or scanning, where applicable) were performed between 9:00 and 12:00 h to minimize effect circadian excursion in corticosterone levels (Gong et al., 2015). Blood sampling was carried out using tail vein bleeding (Kim et al., 2018). The mouse tail was superficially cut with a sterile scalpel blade and blood was collected with EDTA tubes (Microvette® 200 K3 EDTA, Sarstedt, Nümbrecht, Germany). Pressure was gently applied to stop any bleeding before returning to the cage. All blood samples were centrifuged at 4000 g for 5 min at 4°C to obtain plasma, and stored at −80°C until further analysis. Blood corticosterone concentration was measured with enzyme-linked immunosorbent assay (ELISA, Enzo Life Sciences, Inc., USA). All our reported corticosterone concentration levels refer to the amount of corticosterone in plasma, and are expressed in nanograms per milliliter (ng/ml).

#### rsfMRI acquisition

For awake scanning, the mouse was secured (using the headpost) to the custom-made MRI-compatible animal cradle and the body of the mouse was gently restrained by taping its back to the cradle arc as described in the habituation paragraph. For scanning under light anesthesia, mice were first deeply anesthetized with isoflurane (4% induction), intubated and artificial ventilated (90 BPM). In one group, anesthesia was then switched to halothane (0.75%). In a second group, a bolus of medetomidine (0.05 mg/kg) was given via tail vein cannulation before waiting 5 minutes and starting an infusion of medetomidine (0.1 mg/kg/h) with isoflurane reduced to 0.5%. In both cases, the rsfMRI acquisition started 30 minutes after the switch to light anesthesia (Gutierrez-Barragan et al., 2019).

All scans were acquired at the IIT laboratory in Rovereto (Italy) on a 7.0 Tesla MRI scanner (Bruker Biospin, Ettlingen) with a BGA-9 gradient set, a 72 mm birdcage transmit coil, and a four-channel solenoid receive coil (Gutierrez-Barragan et al., 2019). Awake and medetomidine-isoflurane rsfMRI scans were acquired using a single-shot echo planar imaging (EPI) sequence with the following parameters: TR/TE=1000/15 ms, flip angle=60°, matrix=100 x 100, FOV=2.3 x 2.3 cm, 18 coronal slices, slice thickness=550 μm and 1920 time points, for a total time of 32 minutes. Mice under halothane anesthesia (n=19) were scanned with a TR=1200, for a total of 1600 time points, total acquisition time 32 minutes as described in (Gutierrez-Barragan et al., 2019).

### Data analysis

#### rsfMRI data preprocessing

Preprocessing of BOLD images (Fig. S7) was carried out as described in previous work (Gutierrez-Barragan et al., 2019; Grandjean et al., 2020). Briefly, the first 2 minutes of the time series were removed to account for thermal gradient equilibration. RsfMRI timeseries were then time despiked (3dDespike, AFNI), motion corrected (MCFLIRT, FSL), skull stripped (FAST, FSL) and spatially registered (ANTs registration suite) to an in-house mouse brain template with a spatial resolution of 0.2 x 0.2 x 0.6 mm^3^. To optimize awake fMRI timeseries preprocessing we systematically subjected rsfMRI timeseries to a set of increasingly stringent denoising and motion correction strategies involving the regression of either 7, 13 or 25 nuisance parameters (Figure S7A). These were: average cerebral spinal fluid signal plus 6, 12 or 24 motion parameters determined from the 3 translation and rotation parameters estimated during motion correction, their temporal derivatives and corresponding squared regressors (Caballero-Gaudes and Reynolds, 2017). No global signal regression was employed as this procedure artificially introduces regional anticorrelation, and has been reported to reduce antero-posterior extension of fMRI connectivity in the mouse, hence decoupling structural and functional connectivity (Grandjean et al., 2020). In-scanner head motion was quantified via calculations of frame-wise displacement (FD). Average FD levels in awake conditions were comparable to those obtained in anesthetized animals (halothane) under artificial ventilation (p = 0.13, Student t test, Fig. S7B). To rule out a contribution of residual head-motion in awake scans, we further introduced frame-wise fMRI scrubbing using a very stringent FD threshold (0.75 mm, cf. with 1 mm in Grandjean et al., 2020). Quantifications of putatively motion-contaminated volumes at this stringent threshold revealed a predictably higher proportion of labelled frames in awake mice compared to anesthetized subjects (Fig. S7B). However, the use of denoising pipelines of increased stringency, with or without strict frame scrubbing (Fig. S7A), did not reveal any difference in interregional fMRI connectivity strength (p >0.89, one way ANOVA, all regions Fig. S7C), hence arguing against a significant contribution of in-scanner head-motion to our imaging findings. We nonetheless carried out all of our further analyses (both in awake and anesthetized mice) using the most stringent denoising pipeline assessed, i.e. regression of 25 nuisance parameters followed by strict FD > 0.075 mm volume censoring. The resulting time series were band-pass filtered (0.01-0.1 Hz band) and then spatially smoothed with a Gaussian kernel of 0.5 mm full width at half maximum.

#### Seed-based Functional Connectivity

To investigate the topography of functional networks in the awake mouse, we performed a seed-based functional connectivity (FC) investigation using predefined regions of interest (size 4×4×1 voxels) known to be network hubs in the anesthetized mouse brain (Liska et al., 2015; Coletta et al., 2020). These were selected to include both cortical and subcortical systems such as the prefrontal (PFC) and retrosplenial cortex (RS) within the default-mode network (DMN); primary motor (M1) and somatosensory (S1) regions within the latero-cortical network (LCN); visual (VIS) and auditory (A) regions in posterolateral networks (PLN); the insula (Ins) within the salience network (SN); and other subcortical substrates within the hippocampus (HC), thalamus (TH), basal forebrain (BF), and striatum (ST). Canonical correlation mapping was done for each subject by computing the Pearson product-moment correlation coefficient (Pearson’s correlation, for short) between the average signals extracted from each seed, with that of each voxel. The spanned single-subject correlation maps were the transformed using Fisher’s r-to-z transform; averaged across all animals; and thresholded to significant connections with t-scores > 2.8 (two-tailed t-test, p < 0.01, FDR corrected). Averaged maps were then re-transformed to correlation values (r-scores). Group-level seed-based FC differences between awake and anesthetized groups were assessed by means of two tailed, two-sample t-test (p < 0.01, family-wise error cluster-corrected, with defining threshold of t > 2.8).

#### Voxelwise Functional Connectivity Analyses

Voxelwise FC matrices were computed for both awake and anesthetized mice. This was done by computing for each subject the Pearson’s correlation between each voxel and each other voxel in the brain (resulting in a 6843×6843 matrix). Single-subject FC matrices were Fisher transformed and averaged across animals in each group and re-transformed to r-scores. Comparison between groups was done by means of a two-sample t-test (two-tailed, p < 0.05, FDR corrected, critical p =0.009). To avoid topographical bias and help interpretability of our findings, we organized voxels into axonal connectivity modules recently identified in the mouse brain (Coletta et al., 2020). These include a default-mode (DMN); Latero-cortical (LCN), Hippocampus (HC), and Olfactory-Basal Forebrain (OF-BF) networks. Given the observation that some thalamic regions are associated to the axonal DMN module, and others to the LCN, we deliberately separated these into sub-modules for visualization purposes. For the same purposes, we also separated voxels within the DMN corresponding to midline cortico-limbic areas, and posterolateral visual-auditory ones. With this, we computed voxelwise functional connectivity matrices, and averaged them within groups.

#### Structure-function correspondence

Structure-function correspondence was calculated using communities defined by the structural connectome (Coletta et al., 2020). In our prior work we found the identified macro-communities to spatially encompass four macroscopic network systems of the mouse brain that spatially correspond to analogous rsfMRI macro-networks, i.e. the DMN, the LCN, the hippocampus and olfactory-basal forebrain areas (Coletta et al., 2020). To probe the relationship between the functional connectome and the underlying axonal structure we next computed structure-function correspondence (Fig. 4B) as previously described (Cohen and D’Esposito, 2016):

Structure-function correspondence = 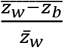

Z_w_ = within-network functional connectivity strength i.e. mean connectivity strength of edges between all pairs of nodes within the same structural network.

Z_b_ = between-network functional connectivity strength i.e. mean connectivity strength of edges between all pairs of nodes that spanned two structural networks.

To depict how intra- and inter-modular communication is distributed across conditions, we mapped the top 10% strongest functional connections at the group level (absolute value, as in (Chelini et al., 2019)) in a network diagram (Figure 4A) using an anatomical parcellation derived from the Allen Brain Institute (Coletta et al., 2020). We further quantified for each subject the mean functional connectivity within voxels within each module, as well as the average of between module connections. We assessed between-group differences with a Mann-Whitney test (p < 0.05, FDR corrected) after retaining for each subject the top 10% strongest connections (Zerbi et al., 2021).

#### Whole brain co-activation patterns (CAPs) analysis

To investigate how rsfMRI dynamics is affected by wakefulness or anesthesia, we used the co-activation patterns approach (Gutierrez-Barragan et al., 2019). This method classifies fMRI volumes into clusters based on their spatial similarity, and the averaged frames within each cluster are then taken to represent recurrent patterns of rsfMRI BOLD co-activation (Gutierrez-Barragan et al., 2019). Specifically, following (Huang et al., 2020), after censoring motion-contaminated frames (FD > 0.075 mm), we ran the k-means clustering algorithm (spatial correlation as distance metric, 500 iterations, 5 replications with different random initializations) on the concatenated data including all frames from all subjects in both awake and anesthetized groups. We set K=6 clusters for two main reasons. First, in previous work (Gutierrez-Barragan et al., 2019) we verified that choosing K = 6 clusters allows to effectively describe most rsfMRI dynamics across multiple halothane-anaesthetized mouse datasets, with over 60% variance explained within each dataset with a limited number of parameters, and diminishing returns using larger cluster numbers; high reproducibility of the clustering algorithm across random initializations; high replicability across independently acquired datasets. Second, we further verified in this study that K = 6 also accounts for a large fraction of variance explained in the collective dataset of awake and anaesthetized mice, with k = 6 being in the “elbow” region of the variance explained as function of number of clusters. The chosen number of clusters was also highly stable across different random initializations, with spatial correlations across CAPs obtained with different initializations greater than 0.99 for all CAPs. For each subject, the fMRI frames classified within a cluster were next voxel-wise averaged to create a single-subject CAP map. Within each group, these maps were then averaged for each CAP at the group level, and thresholded to voxels with significant mean BOLD coactivations or co-deactivations (two-tailed t-test, p < 0.01, FWER cluster-corrected, with defining threshold of t > 2.8). Between-group comparisons for each CAP were done by a voxel-wise two-sample t-test (two-tailed, p < 0.01, FWER cluster-corrected, with defining threshold of t > 2.8).

For each mouse in each group, we computed the occurrence rate of each CAP as the proportion of frames associated to the respective cluster. Between group comparisons of CAP occurrence rates were performed using a two-tailed t-test (p < 0.05, FDR corrected). We then computed the spatial similarity (Pearson’s correlation) between group-averaged CAP maps to verify CAP mirroring motifs structure previously described (Gutierrez-Barragan et al., 2019). We further explored the relation of CAPs to FC by transforming, for each group, the mean co-activation map into a co-fluctuation matrix, multiplying each voxel’s mean BOLD intensity within a CAP with each other voxel’s (Esfahlani et al., 2020). This procedure yields a voxelwise representation of the concordant (or diverging) peaks of BOLD activity that characterize each CAP. Given the robust mirror structure of each CAP anti-CAP pair (cf. Fig. 5B), the resulting cofluctuation matrices are characterized by virtually indistinguishable cofluctuation structure. To spatially link CAP cofluctuation structure to steady-state rsfMRI connectivity, these co-fluctuation matrices were group-averaged and their similarity with the static FC matrix was assessed with Pearson’s correlation.

#### CAP infraslow dynamics

To investigate the dynamics of each CAP, we generated instantaneous CAP-to-frame correlation time courses (Gutierrez-Barragan et al., 2019) at the subject-level. Subsequently, the power spectrum of CAP time courses was computed and averaged for each frequency in each group for the 0.01-0.1 Hz range. To summarize the relative contribution of the infra-slow band to CAP dynamics and that of the Global fMRI Signal (GS), we further computed the fraction amplitude of low frequency fluctuation, or fALFF (Zou et al., 2008), as the proportion of power in the 0.01-0.03 Hz band. fALFF values were compared between groups with a Mann-Whitney test (p < 0.05, FDR corrected). This analysis, and the subsequent investigation of CAP occurrence with respect to Global fMRI Signal (GS) described below, were limited to the sole halothane dataset, as the spectral properties of medetomidine-isoflurane anesthesia present a well-characterized, non-physiological shift towards higher frequencies (Grandjean et al., 2014) that is non representative of the 1/f-like spectral profile observed in awake conditions (cf. Fig. S4).

To map CAP occurrence within the common temporal reference of the phase of the Global fMRI Signal (GS), we computed the instantaneous phase of the filtered (0.01-0.03 Hz) GS using the Hilbert Transform. We collected the GS phase values at each CAP’s occurrence, and to build phase occurrence distributions of CAPs we retained only time frames when the CAP time course at that instant was above 1 SD, in order to ensure we selected only frames reasonably well assigned to a specific CAP (Gutierrez-Barragan et al., 2019). With MATLAB’s CircStats toolbox (Berens, 2009), we tested if the so obtained GS-phase distributions at each CAP’s occurrence deviated from circular uniformity (Rayleigh test, p < 0.001, FDR corrected).

#### Entropy of CAP Markov trajectories

To investigate the dynamics of transitions between CAPs, we defined for each group, a concatenated sequence of CAP occurrences across subjects, and computed the transition probability matrix as the probability of switching from a certain CAP i at time t to another CAP j at time t+1. Only transitions within the same subject were included. We first considered matrices in which we counted the auto-transitions (*i* = *j*), namely persistence probabilities (Cornblath et al., 2020; Huang et al., 2020). The off-diagonal (*i* ≠ *j*), namely transition probabilities from CAP I to CAP j, were then assessed after building matrices from a sequence in which we removed repeating elements in order to control for autocorrelations given the CAP’s dwell time (Cornblath et al., 2020; Huang et al., 2020). The prevalence of a directional transition (*P_ij_* > *P_ji_*) was computed by taking the difference between a transition probability *P_ij_* from *i* to *j* and the transition probability *P_ji_* from *j* to *i*. We used the off-diagonal matrices to measure the entropy of Markov trajectories (HMT) (Ekroot and Cover, 1993; Dimitriadis and Salis, 2017; Huang et al., 2020). This method computes the descriptive complexity of a trajectory between CAPs (in bits), where a higher complexity prescribes more information required to access an ending CAP *j* from an initial CAP *i*; hence it is less accessible as it transitions to other CAPs before reaching its destination. Specifically, for a Markov Chain (MC) representing a CAP sequence with transition probability matrix P, we define the Entropy Rate per step *t* → *t* + 1 as:

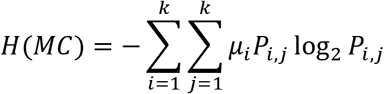

where *μ* is the stationary distribution solving *μ_j_* = ∑*_i_μ_i_P_ij_*. We then define the Markov entropy of a trajectory *T_ij_* from CAP *i* to CAP *j* as:

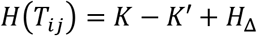

Where *K* = (*I* – *P* – M)^-1^(*H*\* – *H*_Δ_), *I* is the identity matrix, and M is a matrix of stationary probabilities M*_ij_* = *μ_i_*. Here *H*\* is the matrix of single-step entropies *H*\**_ij_* = *H*(*P_i_*) = ∑*_k_μ_k_P_ik_* (from CAP *i* to any CAP *k*), and *H*_Δ_ is a diagonal entropy matrix with trajectories from one CAP to itself (*H*_Δ_)*_ii_* = *H*(*MC*)/*μ_i_*, with zeros if *i* ≠ *j* (Ekroot and Cover, 1993) .

Persistence probabilities were tested for significant deviations from those obtained from random sequences by generating 1000 permutations of CAP occurrence sequences at the subject level and then concatenating the sequences. Differences between each group’s persistence probability was assessed also with these random surrogates and comparing their group differences between auto-transitions with the real one (Huang et al., 2020). Transition probabilities between different CAPs, and the prevalent directional transitions were instead tested for significance above null elements of matrices built after randomly and permuting the non-repeating sequences 1000 times at the subject level before concatenating them in each group (removing an element if *CAP_i_*(*t* + 1) = *CAP_i_*(*t*)). These surrogates were also used to test the significance of between group differences; group-level entropies of Markov trajectories and between group differences.

To rule out the possibility that results were not defined by outlier subjects, we split the groups of subjects into two equal partitions 500 times (Cornblath et al., 2020), and computed the CAP persistence and transition probabilities as well as Markov entropy of the trajectories. Then, we assessed the similarity between the matrices from each split-half sample by means of correlations. We further tested our results at the single-subject level by re-computing for each subject, the persistence and transition probabilities, and comparing them with the results obtained from the concatenated sequences, again by means of correlations between the matrices.

---

**Figure S1.**
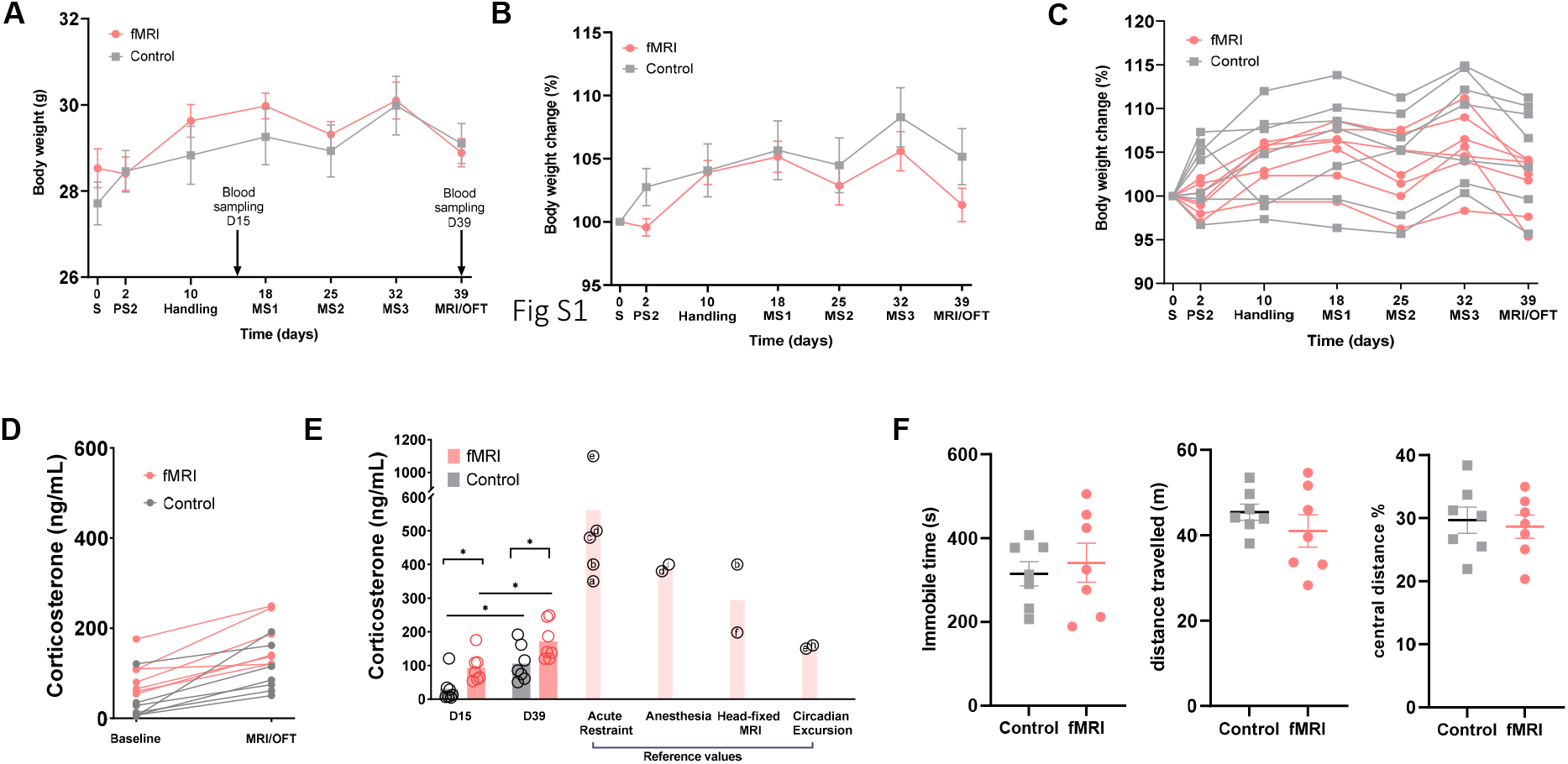
Stress monitoring during the awake habituation procedure. **A)** Body weight measurements over the 4 weeks of habituation protocol (fMRI group, N =7, Control, n = 7). [Day 0 S, before surgery; day 2 PS2, 2 days post-surgery; day 10 Handling, first day of habituation procedure; day 18 MS1, first day of mock scanning session 1; day 25 MS2, first day of mock scanning session 2; day 32 MS3, first day of mock scanning session 3]. Arrows indicate time of blood sampling for corticosterone measurements (D15 and D39). **B)** Percentage change in body weight over the course of the habituation protocol at the group and **C)** subject level. **D)** Plasma corticosterone concentration (ng/mL) at the end of the handling week (fMRI group) or during homecage housing (Control), and right after fMRI scanning (fMRI group) or after open field behavioral testing (control group). **E)** The same values in D) are reported here with respect to representative plasma corticosterone levels reported in the mouse literature upon acute restraint, anesthesia induction, head-fixed MRI. Circadian excursion levels are also reported for reference. [Acute Restraint: a: (Markham et al., 2006), b: (Tsurugizawa et al., 2020), c: (McGill et al., 2006), d: (Snyder et al., 2011), e: (Gong et al., 2015), Anesthesia: d, i: (Kim et al., 2013), Head-fixed MRI: b, f, (Harris et al., 2015), g, (Juczewski et al., 2020); Circadian Excursion: e, h: (Oishi et al., 2006)]. **F)** Time Immobile (seconds, s), distance travelled (meters, m) and percentage of distance travelled in the central zone of the arena in the open field test by mice belonging to the awake fMRI and control littermate groups. Data are expressed as mean ± SEM.

**Figure S2.**
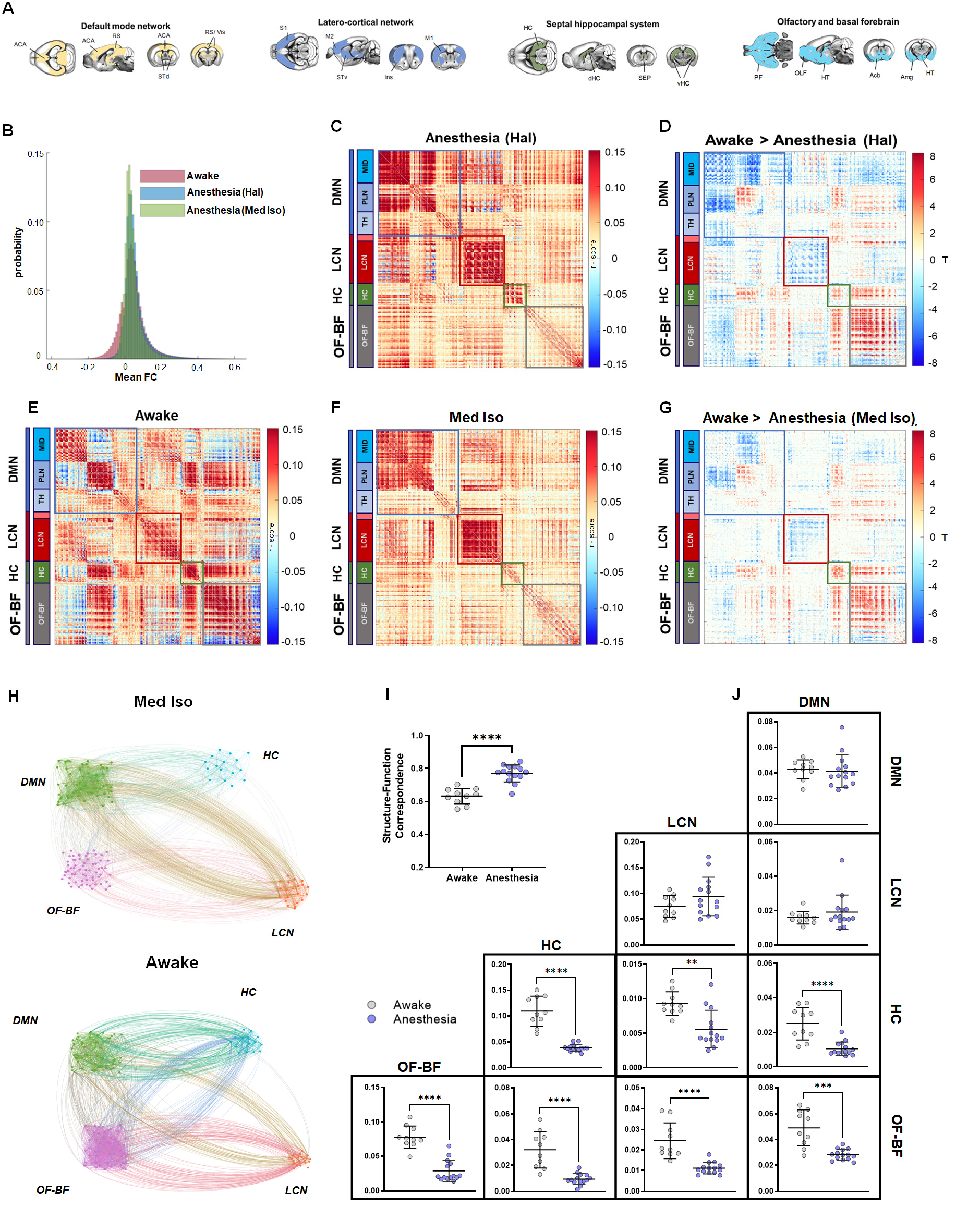
Voxel-vise connectivity and structure-function relationship in medetomidine-isoflurane (Med Iso) anesthetized mice. **A)** Axonal connectivity modules in the mouse brain spatially reconstitute macro-scale networks of the mouse brain (replicated with permission from Coletta et al., 2020). **B)** Distribution of mean functional connectivity in awake and anesthetized mice. Note the larger proportion of negative correlation in awake state. **C-G)** Group averaged voxel-wise FC matrices in awake **(E)** and two independent anesthesia conditions (Halothane and Med Iso, C and F, respectively) with voxels organized within axonal connectivity modules, and further partitioned into network sub-modules. The sub-module corresponding to the thalamic voxels within the LCN module are not labeled (light-red block). Significant between-state FC differences at the voxel level (two-tailed two-sample T-test) for Halothane (D) and Med Iso (G) show consistent deviations from the awake state.**H)** Graphic representation of rsfMRI connectivity within and between previously described axonal modules of the mouse brain (DMN, LCN, HC, OF-BF, from Coletta et al., 2020). Each cluster of nodes represents a subset of anatomically-defined ROIs within the corresponding module. Nodes have been empirically arranged to maximize figure legibility. **I)** Structure-function correspondence in awake and Med Iso anesthetized mice. Between-group differences were assessed with a Mann-Whitney test (p < 0.05). **J)** Quantification of within (diagonal) and between (off-diagonal) network functional connectivity (*p<0.05, ** p<0.01, *** p<0.001, **** p<0.0001, Mann-Whitney test, FDR corrected). [DMN: Default mode network, HC: Hippocampus, OF-BF: Olfactory and basal forebrain, LCN: latero-cortical network].

**Figure S3.**
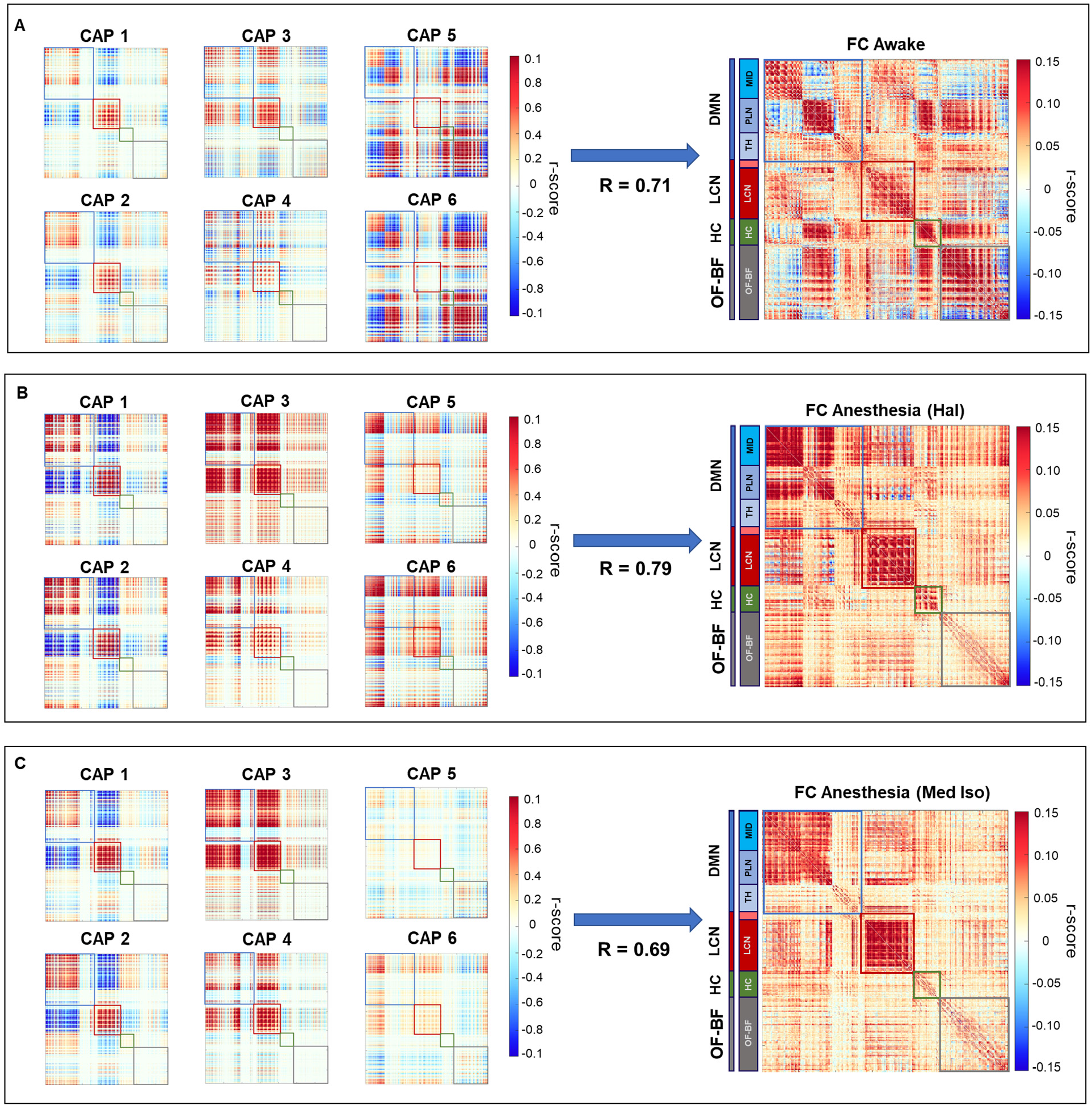
CAPs explain a large fraction of variance in the static functional connectome. Co-fluctuation matrices (left) computed by cross-multiplying each group-mean CAP map with itself, and corresponding time-averaged functional connectivity matrices (right) in awake (top) and anesthetized (Halothane-middle, and Med Iso-bottom) mice. Averaged co-fluctuation matrices (CAPs 1-6) exhibit a correlation of 0.71, 0.79, and 0.69 with the corresponding stationary functional connectome in awake (A), halothane anesthesia (B) and Med Iso, respectively.

**Figure S4.**
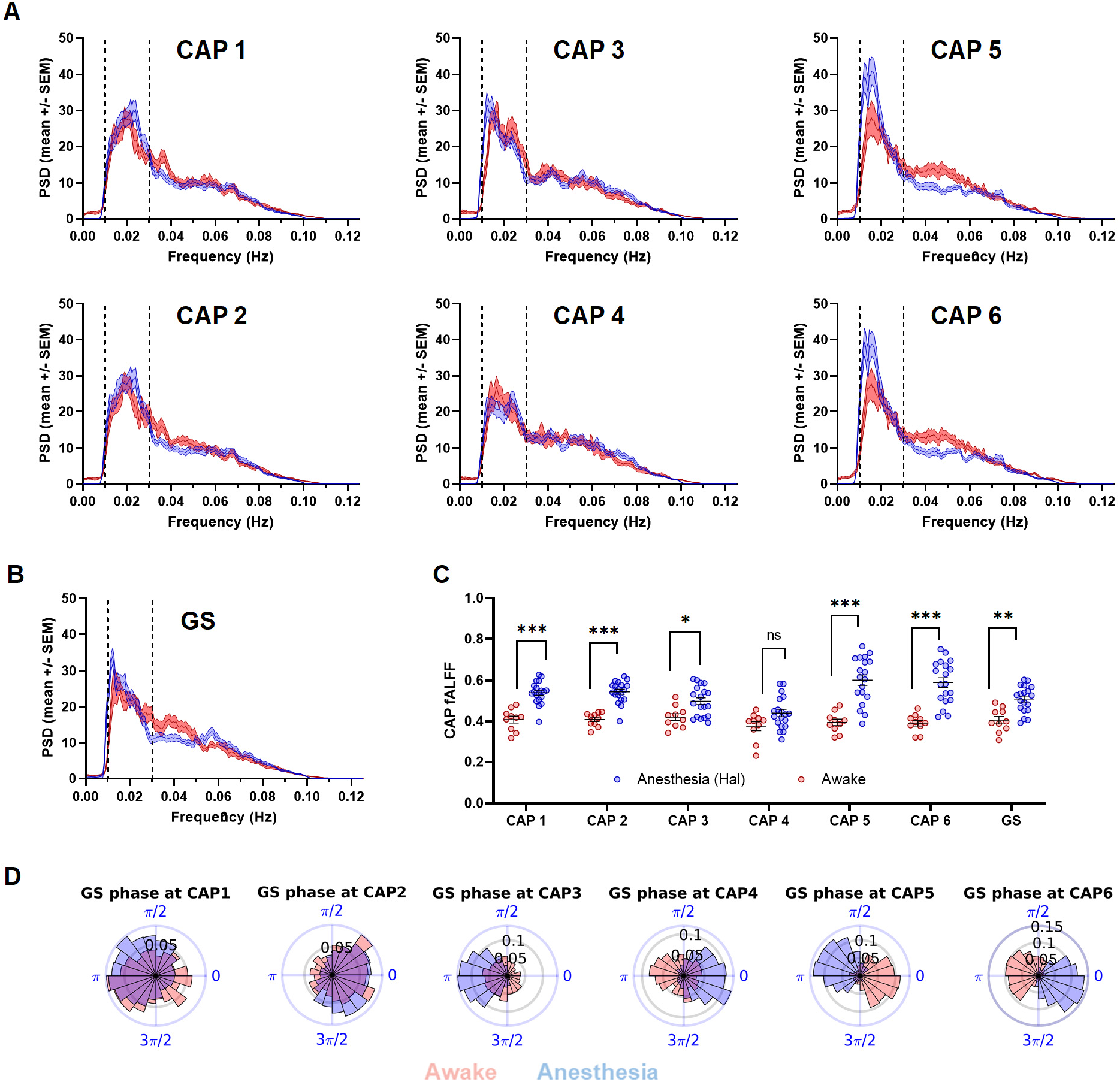
Infraslow dynamics and preferred occurrence of coactivation patterns (CAPs) in awake and anesthetized mice. **A)** Group averaged power spectral density of awake (red) and halothane anesthesia (blue) CAP timecourses (mean +/− SEM). **B)** Power spectral density of the global fMRI signal (GS). **C)** Quantification of the fractional amplitude of low-frequency fluctuations (fALFF) in CAPs and GS spectra, computed as the ratio between 0.01-0.03 Hz band-limited power and the full dynamic range band (0.01-0.1 Hz) for awake (red), and anesthetized mice (blue) (*p<0.05; **p<0.01; *** p<0.001, two-sample T-test, FDR corrected). **D)** Distribution of GS phases at the occurrence of each CAP showing significant deviations from circular uniformity in awake and anaesthetized states (Rayleigh test, p < 0.05 Bonferroni corrected).

**Figure S5.**
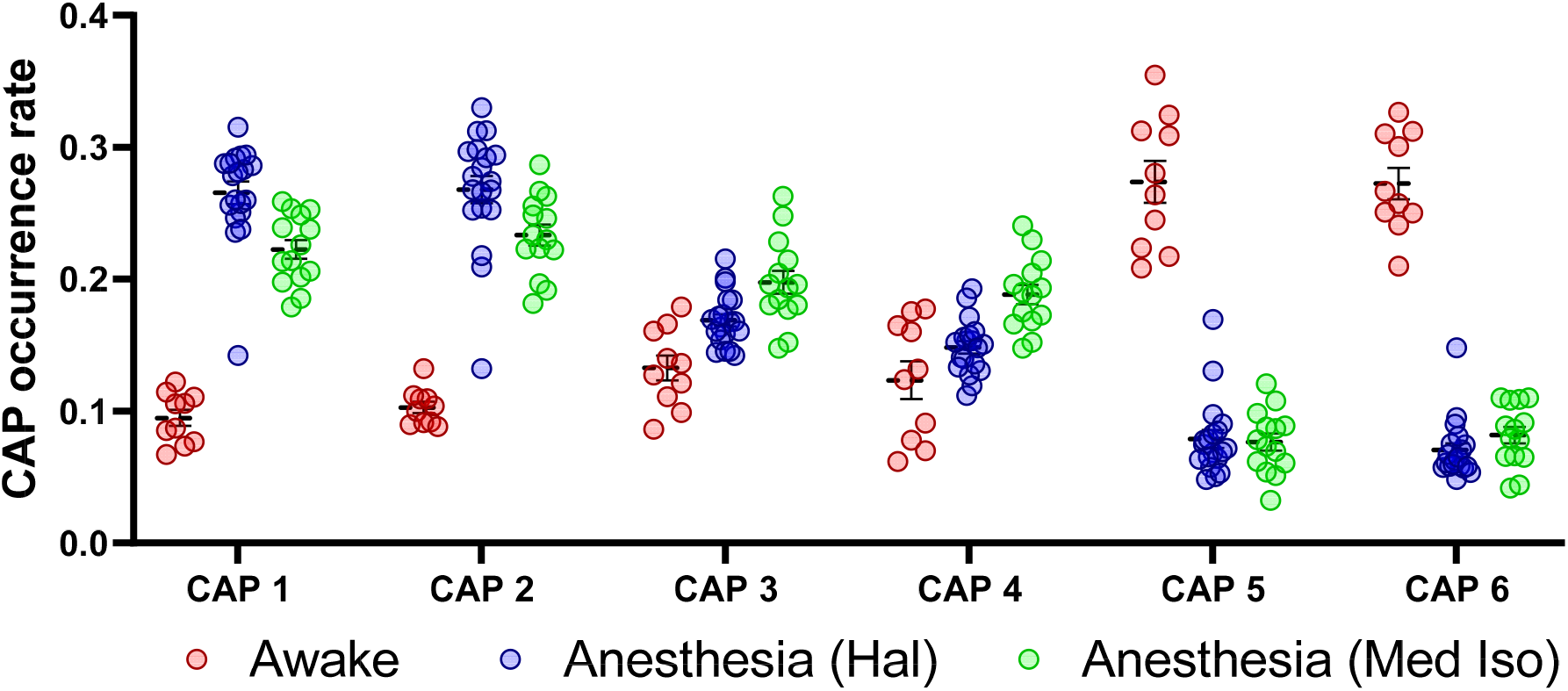
CAP occurrence profile in awake and in anaesthetized animals. Mean (+/− SEM) CAP occurrence rates in wakeful (red), and in halothane (blue) or medetomidine-isoflurane (Med Iso, green) anesthetized mice. CAP occurrence rate with Med Iso anesthesia recapitulates the state-dependent profile observed with halothane.

**Figure S6.**
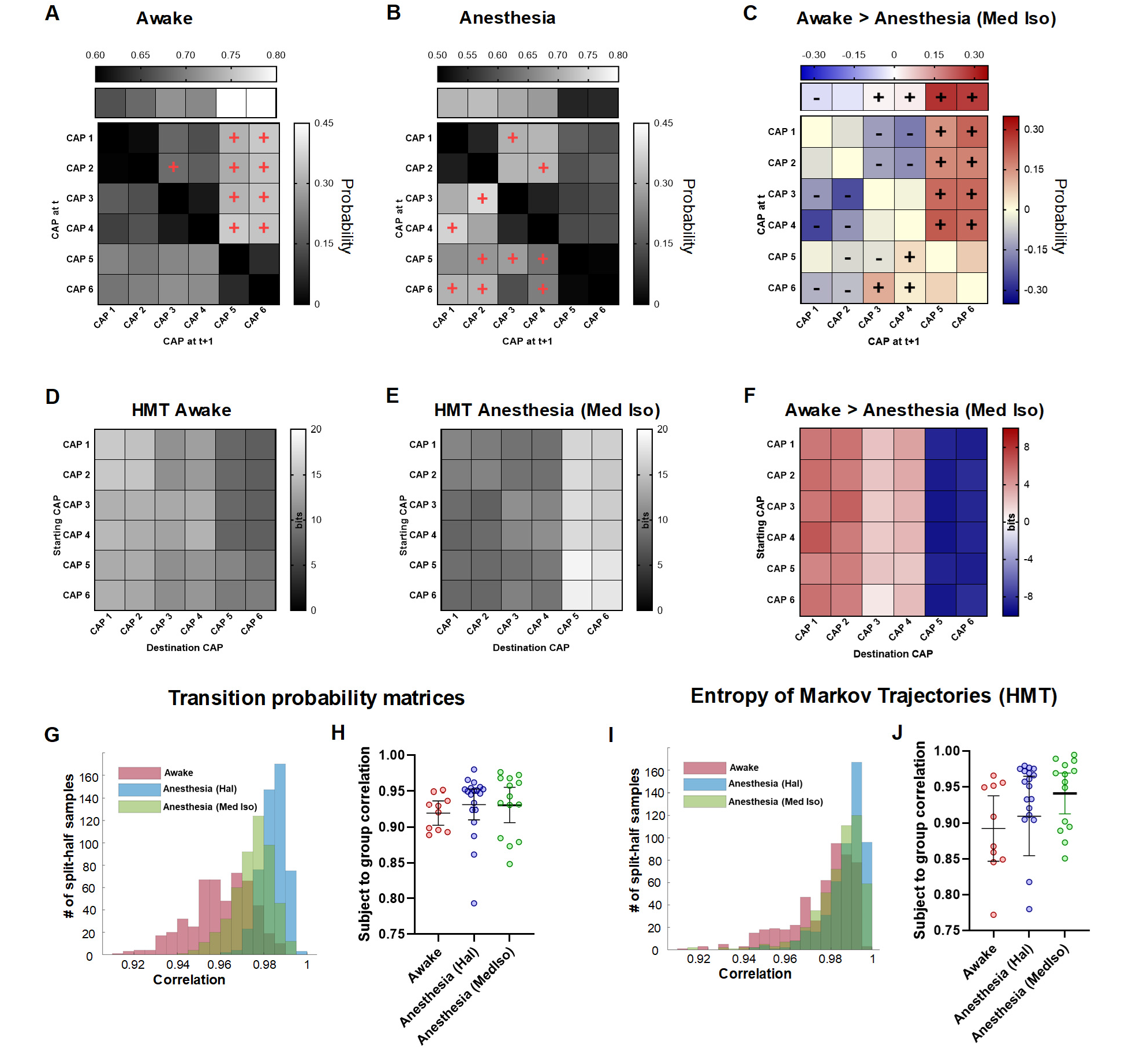
Conserved CAP transition probabilities and preferred trajectories under medetomidine isoflurane anesthesia. **A-B)** Persistence (top row) and transition (off-diagonal) probabilities in awake (A) and Med Iso anesthetized (B) mice. Persistence probabilities are all significant (above null elements from matrices built from 1000 randomly and uniformly permuted sequences not accounting for repetitions). Significant preferred transitions *P_ij_* > *P_ji_* are denoted with a red “+” sign. Directional transitions are depicted if the incoming transition *P_ij_* is significant (above null elements from matrices built from 1000 randomly and uniformly permuted sequences without repetitions). **C)** Transition probability differences between groups are denoted with a “+” sign for transitions higher in awake, and conversely with a “−” sign, if higher in anesthesia. Persistence probability differences were tested with permuted CAP sequences that including repeating elements. **D-E)** Entropy of Markov trajectories (HMT) in awake (D) and anesthetized mice (E). F) Corresponding state-dependent differences in HMT (all elements show significant differences, above and beyond 1000 randomly and uniformly permuted CAP sequences without repetitions). Positive elements represent CAP trajectories with lower accessibility in the awake state as compared with anesthesia, while negative elements are trajectories with facilitated access in awake state. Only the path from CAP 6 to CAP 2 showed non-significant differences. **G)** Distribution of Pearson’s correlation scores between the mean group transition probabilities and the ones obtained after 500 split-half resampling of random subjects within each condition. **H)** Correlations between single subject transition probability matrices and the mean group one. **I)** Distribution of Pearson’s correlation scores between the mean group Entropy of Markov Trajectories matrix and the ones obtained after 500 split-half resampling of random subjects within each condition. **H)** Correlations between single subject Entropy of Markov Trajectories matrices and the mean group one.

**Figure S7.**
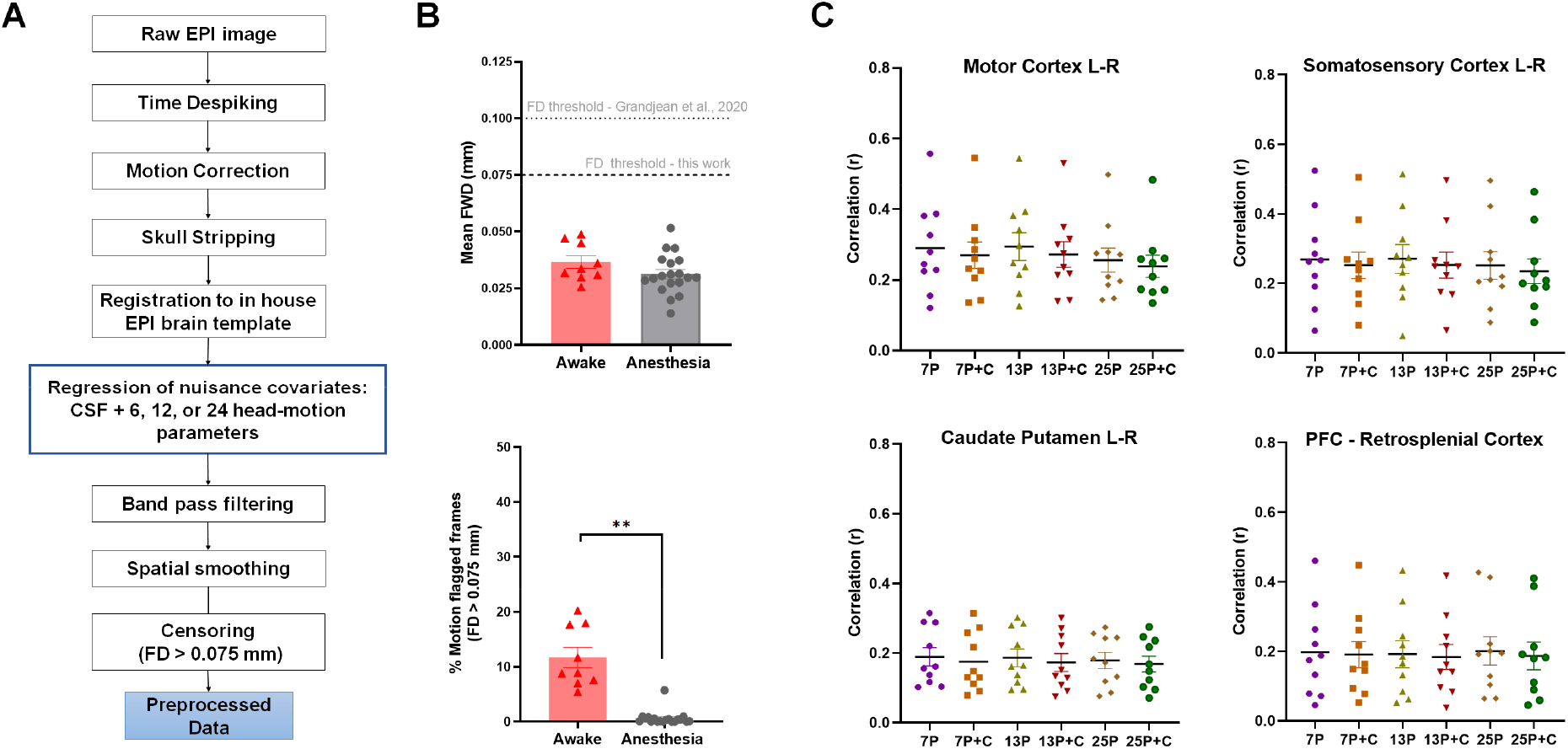
Motion assessment and denoising of awake rsfMRI timeseries. **A)** Preprocessing pipeline used for rsfMRI data denoising. The preprocessing steps investigated in our pilot analyses were the use of increasing head-motion regression parameters (6, 12, or 24), with or without censoring of head-motion flagged frames at a stringent framewise displacement (FD) threshold of 0.075 mm. **B)** Quantification of mean FD (top), and proportion of high FD flagged frames (bottom) in awake and anesthetized (halothane) conditions. **C)** rsfMRI connectivity strength as assessed with a seed-based correlation in awake mice as a function of denoising strategy stringency. As no statistically significant difference in connectivity strength was found across conditions, we opted for the most stringent pipeline assessed (25p + C) for all the analyses of this paper both in awake and anesthetized mice. [7p: regression of CSF and 6 motion parameters; 13P: regression of CSF and 12 motion parameters; 25P: regression of CSF and 24 motion parameters; C: volume censoring].

